# Programmed DNA elimination was present in the last common ancestor of *Caenorhabditis* nematodes

**DOI:** 10.1101/2025.10.23.681605

**Authors:** Lewis Stevens, Simo Sun, Nami Haruta, Yasunobu Maeda, Leyun Xiao, Naoki Uwatoko, Manuela Kieninger, Kazuki Sato, Akemi Yoshida, Dominic Absolon, Joanna Collins, Asako Sugimoto, Taisei Kikuchi, Mark Blaxter

## Abstract

In most organisms, all cells inherit the same genome, and many mechanisms exist to preserve its integrity across cell divisions. Programmed DNA elimination (PDE), the targeted removal of specific genomic regions from somatic cell lineages during early embryogenesis, is a striking exception. Since its discovery in parasitic nematodes over a century ago, PDE has been observed in diverse eukaryotes, including ciliates, arthropods, and vertebrates. However, the mechanisms, functions, and evolutionary origins of PDE remain poorly understood. Here, we describe the discovery of PDE in three species of the free-living nematode genus *Caenorhabditis*. Multiple genomic regions are precisely eliminated from somatic cells during early embryogenesis, resulting in chromosome fragmentation and the loss of key germline genes. The sites of elimination are strongly associated with conserved sequence motifs that likely direct DNA breakage. Comparative analyses indicate that PDE was present in the last common ancestor of *Caenorhabditis* and subsequently lost early during the evolution of many species, including *C. elegans*. The presence of PDE in the ancestors of one of biology’s most important model organisms, together with recent discoveries in other eukaryotic lineages, reveals PDE to be a far more widespread and significant feature of evolution and development than previously recognised.

## Main Text

Early embryonic development is typically characterised by the faithful transmission of identical genomes to all descendant cells. However, some species specifically remove some genomic regions from their somatic cell lineages early during development in a phenomenon known as programmed DNA elimination (PDE). Since its discovery in the late 1800s^1^, PDE has been documented in diverse eukaryotes, including ciliates, nematodes, arthropods, and vertebrates^2^. However, many aspects of PDE, including its evolutionary origins, underlying mechanisms, and biological function, remain obscure^2,3^.

In nematodes, PDE has been most extensively studied in the parasitic family Ascarididae, where it was originally discovered. In ascaridids, DNA is eliminated from all chromosome ends and also from within chromosomes, which results in chromosome fragmentation in somatic cells^4^. The proportion of the genome eliminated in different ascaridid species varies from 18-90%, the majority of which consists of satellite repeats^4–6^. The sites of elimination occur variably within 3-6 kb regions that lack conserved sequence motifs or other identifiable sequence features^4^. Many protein-coding genes are eliminated, some of which have tissue-specific expression in the germline or homologs with known germline functions in related species^5,7^, which has led to the hypothesis that PDE evolved as a mechanism to resolve germline-soma conflict by preventing the ectopic expression of these genes in somatic cells^7^.

PDE was recently discovered in the free-living nematode *Oscheius tipulae*^8^, which is only distantly related to Ascarididae, and far more closely related to the model organism *Caenorhabditis elegans*; both are in suborder Rhabditina. Only a very small proportion of the *O. tipulae* genome (0.6%) is eliminated, and elimination is restricted to the chromosome ends, meaning germline and somatic karyotypes remain the same^8^. Interestingly, the elimination sites in *O. tipulae* are highly precise and contain short palindromic motifs that are required for elimination^9^. PDE has subsequently been described in the free-living rhabditine genus *Mesorhabditis*^10^, suggesting that PDE may be more widespread in nematodes than previously realised.

Here, we report our unexpected discovery of PDE in three early-diverging *Caenorhabditis* species. By combining long-read and chromatin conformation capture sequencing with cytology, we show between 0.7 and 2.3% of the genome is eliminated in each species during early embryogenesis. DNA is eliminated from regions within chromosomes and from chromosome ends, resulting in somatic chromosome fragmentation and the loss of a small number of genes in each species, some of which have *C. elegans* orthologues with essential germline functions. A subset of elimination sites are orthologous across species, and some coincide with genome rearrangement sites. The phylogenetic distribution of PDE suggests that it was present in the last common *Caenorhabditis* ancestor and subsequently lost early during the evolution of many species, including *C. elegans*.

### Programmed DNA elimination in three *Caenorhabditis* species

The *C. elegans* karyotype (five autosomes and one X chromosome) is highly conserved in *Caenorhabditis*^11–19^. As part of an effort to generate chromosome-level reference genomes for all *Caenorhabditis* species, we have sequenced the genomes of 22 species using PacBio HiFi or Oxford Nanopore long-read and chromatin conformation capture (Hi-C) technologies (Table S1). In all cases, the sequenced DNA was extracted from pools of mixed-stage individuals, meaning the majority of cells in each pool were derived from somatic tissues, with only a minority derived from the germline. The Hi-C contact maps for 19 of the 22 species showed six large interacting blocks, consistent with the expected six haploid chromosomes (Fig. 1A; Fig. S1). In contrast, the Hi-C contact maps for *C. auriculariae*, *C. monodelphis,* and *C. parvicauda*, all of which diverge early in the *Caenorhabditis* phylogeny (Fig. 1A), suggested haploid chromosome numbers of 15, 13, and 8, respectively (Fig. 1B-D). Inspection of the ends of these presumptive chromosomes showed that each ended with an array of the nematode telomeric repeat (TTAGGC)n. However, when we aligned PacBio HiFi long reads to these assemblies, we found reads that contained unique sequence extending beyond the telomeric repeat array, albeit at reduced coverage (between 7% and 54% of that of the main chromosome) (Fig. 1H-I). Close inspection of the Hi-C contact maps also revealed interaction between the ends of some of the presumptive chromosomes. This pattern resembles PDE in *O. tipulae*^8^, and suggests that the unexpected assembly fragmentation in *C. auriculariae, C. monodelphis,* and *C. parvicauda* is the result of PDE in these species.

**Fig. 1:**
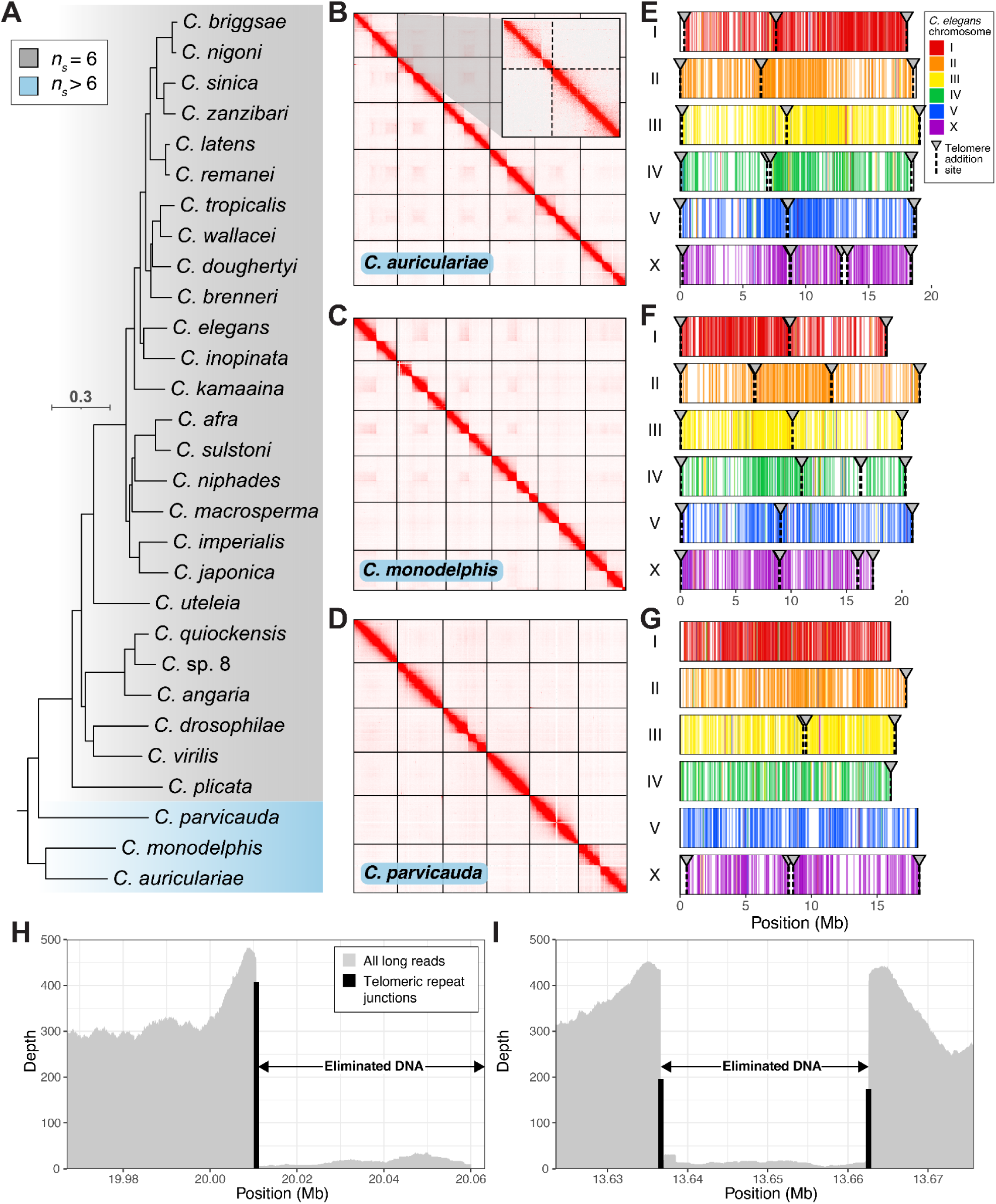
Programmed DNA elimination in three *Caenorhabditis* species. (A) Phylogeny of 29 *Caenorhabditis* species with chromosome-level reference genomes based on 1,601 Benchmarking Using Single Copy Orthologues (BUSCO) nematoda_odb10^20^ genes. Species with haploid somatic chromosome counts (*n_s_*) of 6 are indicated in grey shading and those with counts greater than six (*C. auriculariae*, *C. monodelphis,* and *C. parvicauda*) are highlighted in blue. Branch lengths are in substitutions per site; scale is shown. (B-D) Hi-C contact maps of (B) *C. auriculariae* (inset: map of chromosome I, with dashed lines indicating boundaries of somatic chromosomes), (C) *C. monodelphis,* and (D) *C. parvicauda*. Only the six largest scaffolds are shown. (E-G) Positions of single copy orthologues (from the BUSCO nematoda_odb10 dataset), coloured by the *C. elegans* chromosome (I-V, X) in which the orthologous gene is found, in the six germline chromosomes of (E) *C. auriculariae,* (F) *C. monodelphis,* and (G) *C. parvicauda*. The locations of telomere addition sites are shown with arrowheads and dashed lines. All internally eliminated regions are defined by two telomere addition sites, which can appear as single arrowheads and dashed lines because of the small size of the eliminated regions. (H-I) Depth of all PacBio HiFi reads and of junctions between unique sequence and soft-clipped telomeric repeat in the regions surrounding telomere addition sites on (H) the right end of chromosome III in *C. monodelphis* (III:10,127,500-10,131,500) and (I) the middle of chromosome II in *C. monodelphis* (II:13,623,521-13,675,712). Eliminated DNA is highlighted.

To confirm this, we assembled germline versions of all three genomes (Fig. 1B-D; Table S2). The resulting assemblies for *C. auriculariae* and *C. monodelphis* were highly contiguous (contig N50s of 14.5 Mb and 14.2 Mb, respectively) and nearly all contigs were assigned to chromosomes (100% and 99.6% of bases in the assembly, respectively) (Table S2). In contrast, the *C. parvicauda* assembly was highly fragmented (contig N50 of 0.18 Mb) and had a substantially lower proportion of bases assigned to chromosomes (88.9%), likely due to the high degree of polymorphism present in the sequenced strain (Fig. S2). Despite these differences, all three germline genome assemblies contained six chromosome-sized scaffolds that were clearly homologous to the six *C. elegans* chromosomes (I-V, X) (Fig. 1E-G).

To identify eliminated regions in the three *Caenorhabditis* species, we aligned PacBio HiFi reads to each genome and searched for instances of telomeric repeat-containing reads mapping within the germline chromosomes (Fig. 1H-I). In *C. auriculariae*, we detected sites of telomeric repeat array addition (hereafter referred to as telomere addition sites) at positions near both ends of all six chromosomes (Fig. 1E). We also identified pairs of telomere addition sites that defined seven internally eliminated regions, suggesting that the six germline chromosomes are fragmented into 13 somatic chromosomes (*n_g_*=6, *n_s_*=13). In *C. monodelphis*, we identified eliminated regions at all chromosome ends and at nine internal regions, generating 15 somatic chromosomes (*n_g_*=6, *n_s_*=15) (Fig. 1F). In *C. parvicauda*, we identified two internally eliminated regions, generating eight somatic chromosomes (*n_g_*=6, *n_s_*=8), and identified eliminated regions at 5 of the 12 germline chromosome ends (Fig. 1G). We do not know whether the remaining ends undergo elimination because they are unresolved in the assembly. We did, however, identify one additional eliminated region in a short scaffold that we could not associate with any of the chromosome-sized scaffolds, which is likely derived from an incomplete chromosome end. Across the three species, the eliminated regions ranged in size from 1.3 kb to 500.6 kb. In total, we inferred that at least 2.53 Mb (2.3%), 0.88 Mb (0.7%), and 1.31 Mb (1.1%) of DNA is eliminated from the genomes of *C. auriculariae, C. monodelphis*, and *C. parvicauda*, respectively (Table S3).

### Sequence motifs at telomere addition sites

In *O. tipulae,* the telomere addition sites are precise and contain palindromic motifs required for PDE^9^. With the exception of two sites in *C. monodelphis*, where telomere addition occurred heterogeneously within 1-2 kb regions containing satellite repeat arrays (Fig. S3), the telomere addition sites in the three *Caenorhabditis* species were also precise (Fig. 1H-I), with less than 2% of telomeric repeat-containing reads supporting alternative positions.

To identify sequence motifs associated with the telomere addition sites, we used MEME^21^ to search the regions surrounding each site for overrepresented sequences. In *C. auriculariae*, we identified a pair of motifs that were present at all telomere addition sites (except at one end of chromosome V, where the assembly is incomplete due to the presence of the ribosomal DNA repeat array). The motifs (motif-1 and motif-2) were 14 bp and 11 bp in length, respectively, and were separated by a gap ranging from 9 to 50 bp (Fig. 2A; Figure S4). We refer to this pair of motifs as the *C. auriculariae* Sequence For Elimination (Ca-SFE). The Ca-SFE typically occurred upstream of the telomere addition site, within the retained DNA (Fig. S4-5; Data S1). While the pair of motifs typically occurred in the same orientation, we identified four sites where the first motif was inverted (Data S1). In *C. monodelphis*, we identified an SFE (Cm-SFE) that was similar in sequence and structure in all 28 of the precise telomere addition sites (Fig. 2A-B; Fig. S4). The motifs were 13 bp and 11 bp in length, respectively, and were separated by a gap of between 15 and 68 bp. In *C. parvicauda*, we identified a SFE that differed in both sequence and structure from those in *C. auriculariae* and *C. monodelphis*. The Cp-SFE was 24 bp, non-palindromic, and lacked an internal gap (Fig. 2A). It was present at all 10 fully assembled telomere addition sites, occurring between 27 and 34 bp upstream (Fig. S4-5; Data S1). None of the *Caenorhabditis* SFEs shared sequence similarity to the *O. tipulae* SFE.

**Fig. 2:**
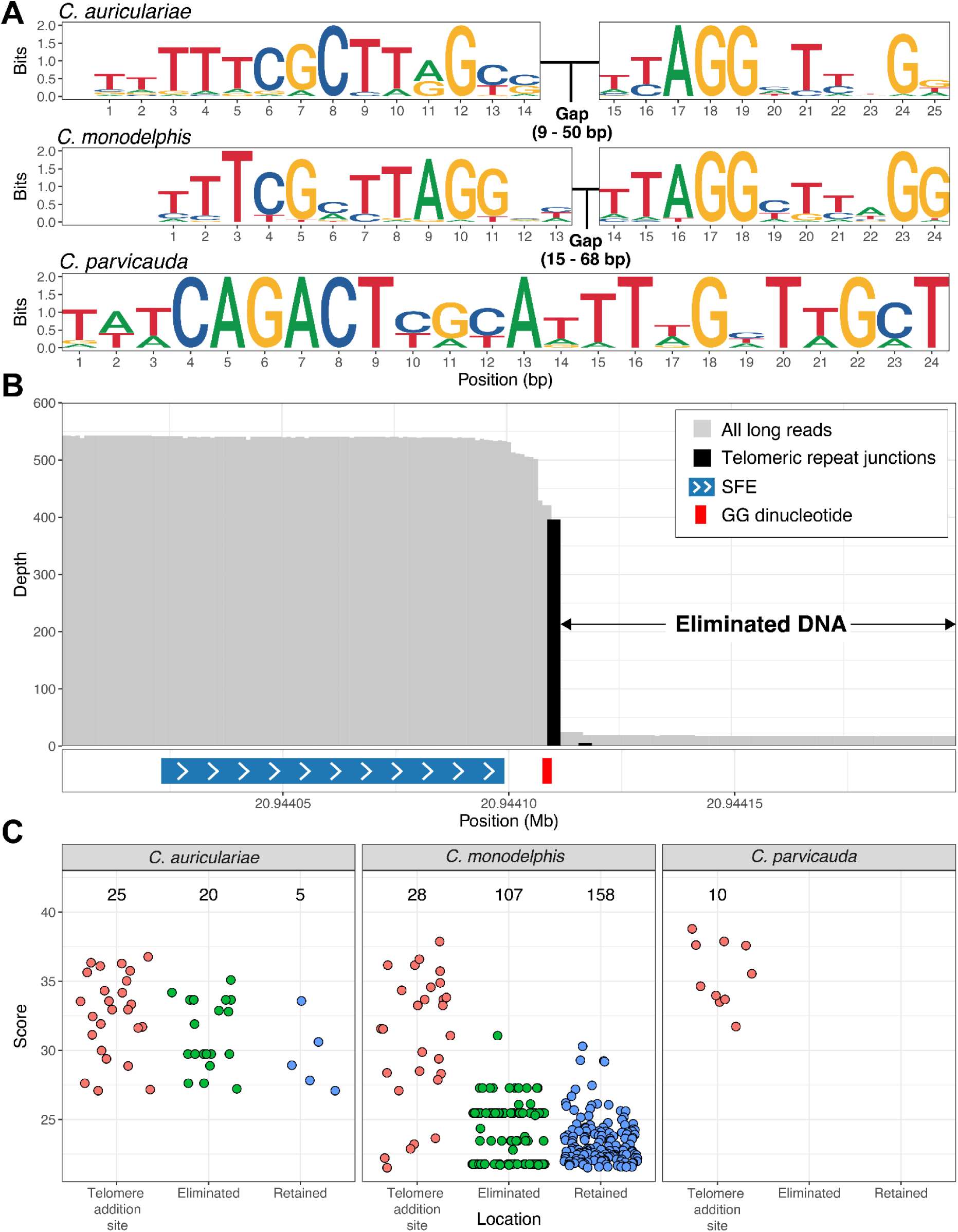
Sequence motifs are present at telomere addition sites. (A) Sequence logos of the sequences for elimination (SFEs) associated with the telomere addition sites in *C. auriculariae*, *C. monodelphis*, and *C. parvicauda*. The *C. auriculariae* and *C. monodelphis* SFEs are aligned to highlight their similarity. The ranges of gap lengths between the *C. auriculariae* and *C. monodelphis* SFEs are indicated. (B) The position of the SFE relative to the telomere addition at the right end of V in *C. monodelphis* (V:20,944,000-20,944,200). Depth of all PacBio HiFi reads and of junctions between unique sequence and soft-clipped telomeric repeat are shown. Position of GG dinucleotide is shown. (C) Scores of SFE occurrences associated with telomere addition, eliminated DNA, and retained DNA. Numbers indicate the number of SFE hits. Only SFE occurrences that had scores equal to or greater than the lowest scoring telomere addition site-associated occurrences (27.1 in *C. auriculariae*, 21.5 in *C. monodelphis*, and 33.5 in *C. parvicauda*) are shown. SFEs that occurred within the germline telomeric repeat blocks at the end of each chromosome are not shown.

To test the specificity of the SFEs, we scanned each genome using a position-weight matrix derived from alignments of the corresponding SFE sequences. The Cp-SFE was highly specific, with all high-scoring matches being associated with telomere addition sites (Fig. 2C). In contrast, while the majority of highest-scoring Ca-SFE and Cm-SFE matches were associated with telomere addition sites, we also found many high-scoring matches elsewhere (Fig. 2C). This included 20 (*C. auriculariae*) and 107 (*C. monodelphis*) matches within eliminated DNA that scored at least as high as the lowest-scoring SFE linked to telomere addition sites. Although we found no evidence of telomere addition at these sites, there is evidence in *O. tipulae* that some alternative SFEs undergo breakage and telomere addition, but these events are too rare to be captured by standard genome sequencing experiments^9^. In the retained DNA, 5 (*C. auriculariae*) and 158 (*C. monodelphis*) matches scored above the same threshold, although most were relatively low-scoring (Fig. 2C).

Close inspection of somatic and germline chromosome ends also revealed several patterns related to telomere maintenance in the three species. First, the telomeric repeat arrays differed substantially in length between the germline and soma. In *C. auriculariae* and *C. monodelphis*, somatic chromosome ends had a median telomere lengths of 354 bp and 768 bp, respectively, whereas germline chromosome ends had lengths of 3.1 kb and 10.2 kb (Fig. S6A). Although direct comparison of germline and somatic chromosome ends in *C. parvicauda* was not possible due to the fragmented genome assembly, a subset of telomeric repeat-containing reads contained very long arrays (>4 kb) (Fig. S6B), suggesting germline telomeres are also longer in *C. parvicauda*. Second, in *C. auriculariae* and *C. monodelphis*, somatic telomere addition sites were consistently preceded by a ‘GG’ dinucleotide (Fig. 2B; Fig. S5A; Fig. S7A-B), which may serve to initiate telomeric repeat array elongation. This pattern was not present in *C. parvicauda* (Fig. S7C). Finally, in *C. auriculariae*, germline telomeric repeat array sequences showed periodic patterns at four chromosome ends (Fig. S8), consistent with activity of an alternative lengthening of telomeres (ALT) mechanism, which has been reported in telomerase-deficient mutants of *C. elegans*^22^.

Together, our results show that PDE-associated telomere addition sites in *Caenorhabditis* are highly precise and strongly associated with conserved sequence motifs that likely serve as binding sites for the as-yet-unidentified elimination machinery.

### A subset of eliminated genes have known germline functions in *C. elegans*

PDE in nematodes has been hypothesised to be a mechanism for avoidance of germline-soma conflict by preventing genes with germline-specific functions from being expressed in the soma^4,5,7^. To identify eliminated genes in the three *Caenorhabditis* genomes, we predicted protein-coding genes in the new genomes and clustered them into orthologous groups along with the gene sets from other *Caenorhabditis* species. We identified 196 (1.2% of the total gene set), 81 (0.5%), and 115 (0.7%) eliminated genes in *C. auriculariae*, *C. monodelphis*, and *C. parvicauda*, respectively. Most of these eliminated genes were present in multiple copies in the genome (Fig. 3A; Data S2). For example, 82% (161/196) of the eliminated genes in *C. auriculariae* were multi-copy, with 53% (103/196) having at least one additional copy in the retained portion of the genome (Fig. 3A). The majority of eliminated genes also lacked *C. elegans* homologs: 80% (*C. auriculariae*), 65% (*C. monodelphis*), and 69% (*C. parvicauda*) of the eliminated genes belonged to orthologous groups without a *C. elegans* representative (Data S2). Although we found 12 orthogroups with members that were eliminated in two or more of the three eliminating species, the majority of these were present in multiple copies in the genome and comprised transposon-derived proteins (Data S2).

**Fig. 3:**
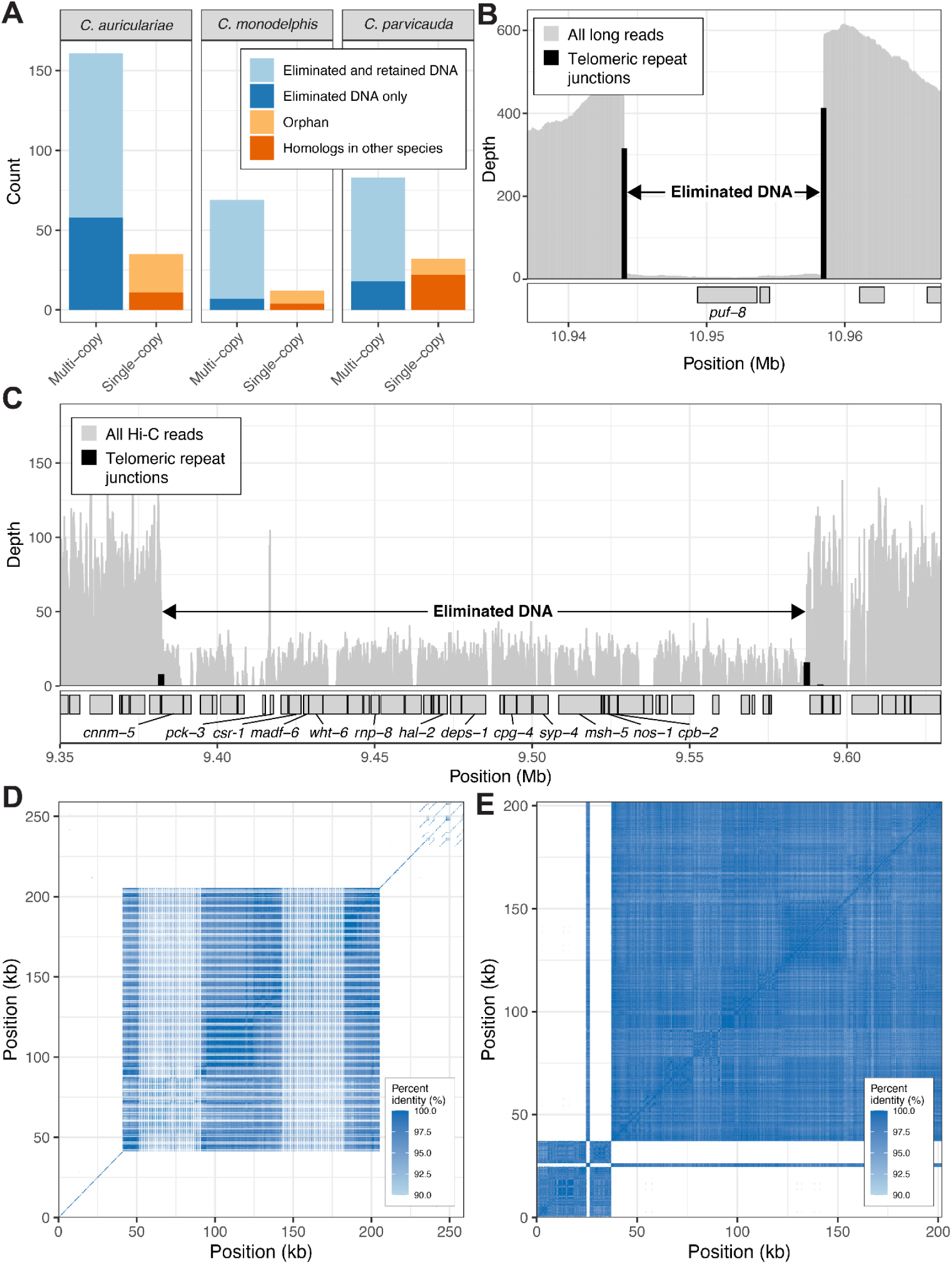
Eliminated DNA contains germline-associated genes and satellite repeats. (A) Summary of eliminated genes in each species based on orthology clustering alongside proteins from related species. ‘Eliminated and retained DNA’ refers to eliminated genes belonging to orthogroups with eliminated and non-eliminated members, whereas ‘Eliminated DNA only’ refers to eliminated genes belonging to orthogroups with eliminated members only. ‘Orphan’ refers to genes belonging to orthogroups without members from other species, whereas ‘Homologs in other species’ refers to genes belonging to orthogroups without members from other species. (B) Depth of all PacBio HiFi reads and of junctions between unique sequence and soft-clipped telomeric repeat across an eliminated region on chromosome IV containing *puf-8* (IV:10.937-10.967 Mb). Eliminated DNA is highlighted. (C) Depth of all Hi-C reads and of junctions between unique sequence and soft-clipped telomeric repeat across an eliminated region on chromosome III in *C. parvicauda* (III: 9.32-9.65 Mb). Eliminated DNA is highlighted. Hi-C read depth is shown because PacBio HiFi read depth is highly variable, owing to polymorphism in the strain we sequenced (Fig. S2). Genes that are one-to-one orthologues of *C. elegans* germline genes are labeled. (D) Self-identity heatmap of the 259 kb region eliminated from the middle of chromosome IV in *C. auriculariae* containing 163 kb of a 150 bp repeating unit (Ca_SR8). (E) Self-identity heatmap of the 202 kb region eliminated from the right end of chromosome IV containing 165 kb of a 160 bp repeating unit (Ca_SR2).

We did, however, find a small number of eliminated genes in each species that were one-to-one orthologues of *C. elegans* genes with known germline-specific functions. In both *C. auriculariae* and *C. monodelphis*, the orthologue of *puf-8* is eliminated (Fig. 3B; Fig. S9; Data S2). PUF-8 is a Pumilio-family protein that binds to the 3’-UTR of germline mRNAs and has multiple roles in the *C. elegans* germline, including controlling germline stem cell proliferation and differentiation^23–25^. These *puf-8* orthologues are encoded on different chromosomes in the two species: the *C. monodelphis puf-8* orthologue is encoded in a 14.4 kb internally eliminated region of chromosome IV (Fig. 3B), whereas the *C. auriculariae* orthologue is found in a 73.6 kb internally eliminated region of chromosome V (Fig. S9).

In *C. parvicauda*, 17 of the 115 eliminated genes have one-to-one orthologues in *C. elegans*, and all were encoded on a single internally eliminated region on chromosome III (Fig. 3C). Several of these genes encode RNA-binding proteins whose orthologues have known germline functions in *C. elegans*. For example, *rnp-8* is eliminated in *C. parvicauda*. In *C. elegans*, RNP-8 is a component of a cytoplasmic poly(A) polymerase that promotes oogenesis^26^. Another eliminated gene, *cpb-2*, encodes an RNA-binding protein responsible for cytoplasmic polyadenylation, which regulates the translation of mRNAs in the germline^27^. Similarly, *nos-1*, a member of the Nanos family, is known to regulate primordial germ cell development^28^. Other eliminated genes are involved in other aspects of germline function, such as *deps-1*, a key component of the P-granule assembly pathway^29^, and *hal-2*, *syp-4*, and *msh-5*, which play critical roles in meiosis, with *hal-2* promoting homologous chromosome pairing^30^, *syp-4* being a component of the synaptonemal complex^31^, and *msh-5* promoting crossover formation^32^. Lastly, the orthologue of *csr-1* is eliminated, which encodes an Argonaute protein with multiple germline roles, including regulating gene expression and promoting chromosome segregation^33–35^.

### Satellite repeats and helitrons are common in eliminated DNA

Satellite repeats make up a substantial proportion of the eliminated DNA in both ascaridid and *Mesorhabditis* nematodes^4–6,10^, but only a small proportion in *O. tipulae*^8^. We identified satellite repeats in the genomes of all three species and found that they were significantly enriched in eliminated DNA (*P* < 0.05 in all three species) (Fig. S10; Table S4). In *C. auriculariae*, where satellite repeats make up 1.5% of the whole genome, 26% of the eliminated DNA is composed of satellite repeats. For example, in one 259 kb internally eliminated region on chromosome IV, 163 kb (63%) is composed of a 150 bp tandemly-repeating unit (Ca_SR8; Fig. 3D). Similarly, 165 kb (82%) of a 202 kb region eliminated from the right end of chromosome IV is composed of a 160 bp tandemly-repeating unit (Ca_SR2; Fig. 3E). Although most eliminated regions contained distinct repeat units (Fig. S10), we found cases where the same repeat unit appeared in multiple separate eliminated regions in a species. For example, the Ca_SR2 repeating unit found in the right end of *C. auriculariae* chromosome IV also makes up 80% and 69% of the DNA eliminated from the right end of chromosome I and left end of chromosome III, respectively (Fig. S10). No satellite repeat units were shared across the three species.

We next asked whether transposable elements (TEs) were also enriched in eliminated DNA. Overall, TEs were underrepresented in eliminated DNA (Fig. S11), likely because of the high satellite repeat content. However, certain families were significantly enriched, such as the DNA-TCMar-m44 family in *C. auriculariae* (Fig. S11). Interestingly, we found that helitrons, which are present in eliminated DNA in *O. tipulae*^8,9^, were significantly enriched in eliminated DNA in *C. auriculariae* and *C. monodelphis* (*P* < 0.001 in both species). The vast majority of these were located in the eliminated regions at chromosome ends (100% and 80%, respectively). However, we also found that helitrons were significantly enriched (*P* < 0.05) in the chromosome ends of 5 of the 26 non-eliminating *Caenorhabditis* species with chromosome-level reference genomes, including *C. elegans*, suggesting that the enrichment of helitrons in eliminated DNA may reflect a preferential association with chromosome ends generally, rather than with eliminated DNA specifically.

### Timing and cytology of PDE in *C. auriculariae*

To investigate the cytology of PDE in *Caenorhabditis*, we focused on *C. auriculariae,* which eliminates the largest amount of DNA among the three species. We analysed early embryonic cell divisions using DIC microscopy, which revealed that cleavage patterns were largely conserved between *C. elegans* and *C. auriculariae*, although the latter had a longer cell cycle and the P4 blastomere divided before the C blastomere rather than after (Fig. 4A). Hereafter, the cell nomenclature of *C. elegans* is used for *C. auriculariae*.

**Fig. 4:**
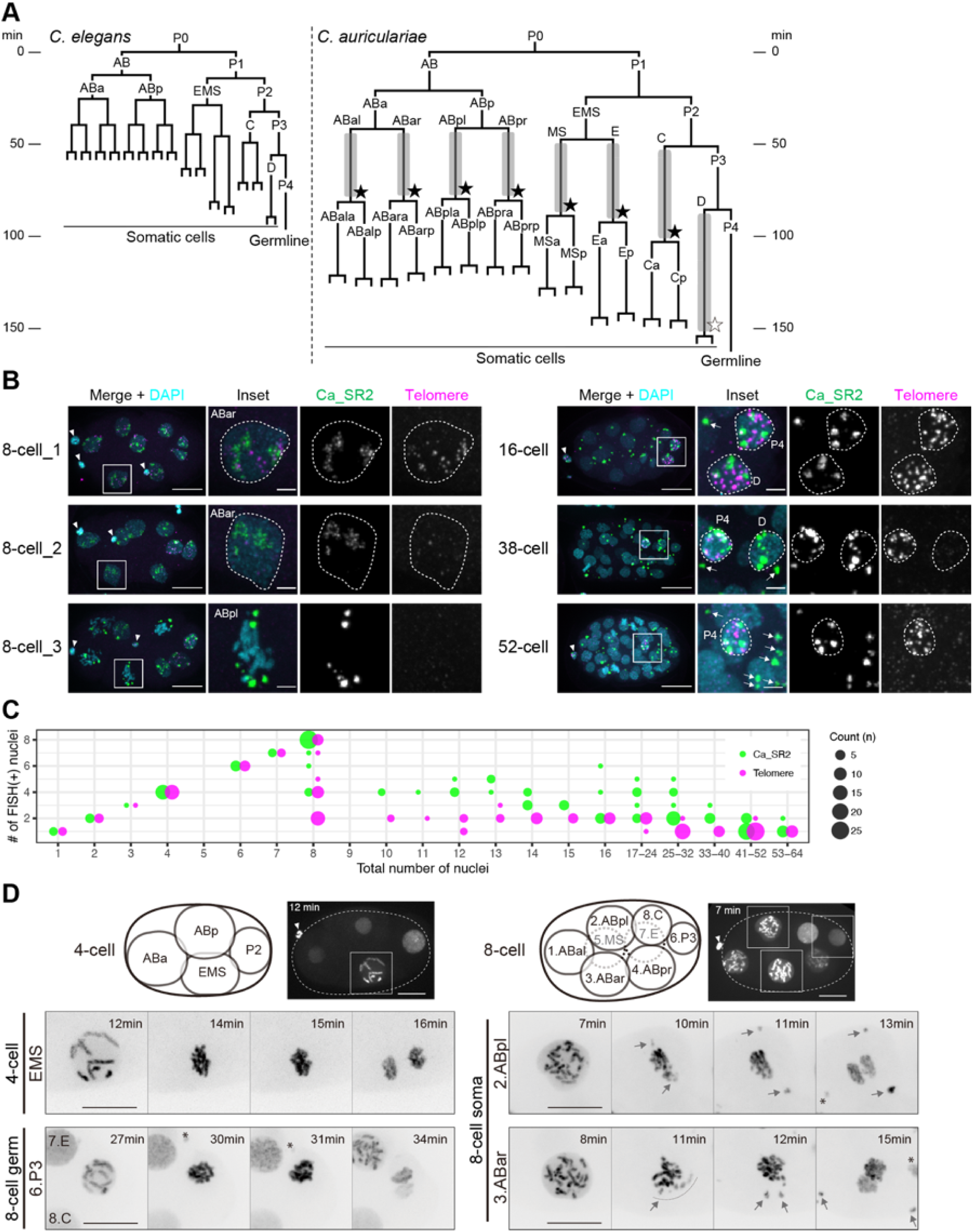
DNA elimination in *C. auriculariae* embryos. (A) Early embryonic cell lineage of *C. elegans* and *C. auriculariae*. Timeline indicates minutes after the first cleavage. Grey bars, periods of telomere signal reduction; black stars, mitosis when eliminated DNA fragments are excluded from nuclei; open star, inferred timing of DNA elimination in the D cell based on the Ca_SR2 and Ca_SR8 signals. (B) FISH of early embryos with probes for eliminated sequences. Ca_SR2 satellite repeats (green) and telomeric repeats (magenta), with DNA counterstained by DAPI (cyan). Insets show magnified views of boxed regions. 8-cell_1, ABar early interphase; 8-cell_2, ABar late interphase; 8-cell_3, ABpl metaphase; 16-cell, D early interphase and P4 interphase; 38-cell, D late interphase and P4 interphase; 52-cell, P4 interphase. Arrows, cytoplasmic DNA fragments; arrowheads, polar bodies. Scale bars: 10 μm (embryos), 2 μm (insets). (C) Quantification of nuclei with Ca_SR2 and telomere foci (n = 169 embryos). X-axis: total nuclei count per embryo (DAPI). Y-axis: nuclei with FISH signals comparable to germline cells. Dot size indicates number of embryos assayed. (D) Time-lapse confocal images of GFP::Histone H2B during mitosis in 4- and 8-cell embryos. (Top) Schematics of blastomere positions and whole-embryo images. (Bottom) Magnified views of indicated nuclei with inverted colors. Arrowheads, polar bodies; grey arrows/dotted lines, eliminated DNA fragments; asterisks, signals from adjacent cells. Scale bars: 10 μm.

To determine the timing and cell lineage occurrence of PDE in *C. auriculariae*, we performed DNA fluorescence in situ hybridization (FISH) using probes for the telomeric repeat (TTAGGC)n (expected to be found at all chromosome ends) and Ca_SR2 satellite repeats, which are confined to the terminally eliminated regions of chromosome I (right), III (left), and IV (right) (Fig. 4B,C; Fig. S10). All nuclei showed both signals before the 8-cell stage (Fig. 4C). During 8-cell interphase, Ca_SR2 signals remained unchanged in all cells, whereas telomeric repeat signals significantly weakened in all somatic nuclei but persisted in the germline precursor P3 (Fig. 4B; 8-cell_1, 8-cell_2). During somatic mitosis at the 8-cell stage, Ca_SR2 signals were detected at the periphery of the metaphase plate (Fig. 4B; 8-cell_3), and were later observed in the cytoplasm after nuclear reformation (Fig. 4B; 16-cell, 38-cell, 52-cell). In the D cell (somatic daughter of P3), telomeric repeat signals also weakened during interphase, whereas Ca_SR2 signals persisted (Fig. 4B; 16-cell, 38-cell). After D cell division around the 40-50-cell stage (Fig. 4A; 150 min), telomeric repeat and Ca_SR2 signals were detected only in one or two cells, presumed to be germ lineage cells (P4, or its daughters Z2 and Z3) (Fig. 4B; 52-cell). Cytoplasmic Ca_SR2 signals persisted for several more divisions but became undetectable by the 200-cell stage (Fig. S12A; 100-200-cell), suggesting the degradation of eliminated DNA. Ca_SR8, a satellite repeat located in the internal eliminated region of chromosome IV, exhibited spatiotemporal patterns similar to Ca_SR2 (Fig. S12). The telomeric repeat signals in somatic cells remained weaker than those in germline cells in later embryonic stages (Fig. S13), consistent with our finding that somatic telomeric repeat arrays were much shorter than germline telomeres (Fig. S7).

We further analysed PDE by live-imaging of GFP::Histone H2B transgenic nematodes (Fig. 4D; Movies S1-S2). In 8-cell embryos, condensed chromosomes in somatic blastomeres appeared shorter and more numerous than those at the 4-cell stage (Fig. 4D; 7 min ABpl and 8 min ABar). In somatic blastomeres, small chromosome fragments were detected at the periphery of the metaphase plate (Fig. 4D; 10 min ABpl and 11 min ABar), which rapidly moved toward the cell cortex before anaphase, and were excluded from the reforming nucleus. Consistent with FISH, no chromosomal fragmentation or eliminated DNA was observed in the P3 cell (Fig. 4D).

Taken together, PDE in *C. auriculariae* occurs in all somatic lineages during early embryogenesis, with telomere shortening preceding the exclusion of eliminated chromosome fragments from nuclei (Fig. 4A).

### Some elimination sites are orthologous and some coincide with rearrangement sites

We next sought to determine whether any sites of elimination were orthologous between the three species. To do this, we compared the relative positions of orthologous genes in the *C. auriculariae, C. monodelphis*, and *C. parvicauda* genomes and overlaid the positions of eliminated DNA (Fig. 5; Fig. S14). Consistent with previous observations in *Caenorhabditis*^14–17,19,36,37^, the vast majority of genes have remained on the same chromosome, but their order has been highly rearranged (Fig. 5; Fig. S14). Gene order was more conserved on the X chromosomes than autosomes (Fig. 5; Fig. S14). We observed a large synteny break in the X chromosomes of *C. auriculariae* and *C. monodelphis*, which was consistent with two large inversions each affecting approximately half of the chromosome (Fig. 5). This break coincided with an internally eliminated region (Fig. 5): the two genes that flank this region in the *C. auriculariae* X chromosome are found at opposite ends of the *C. monodelphis* X, and vice versa.

**Fig. 5:**
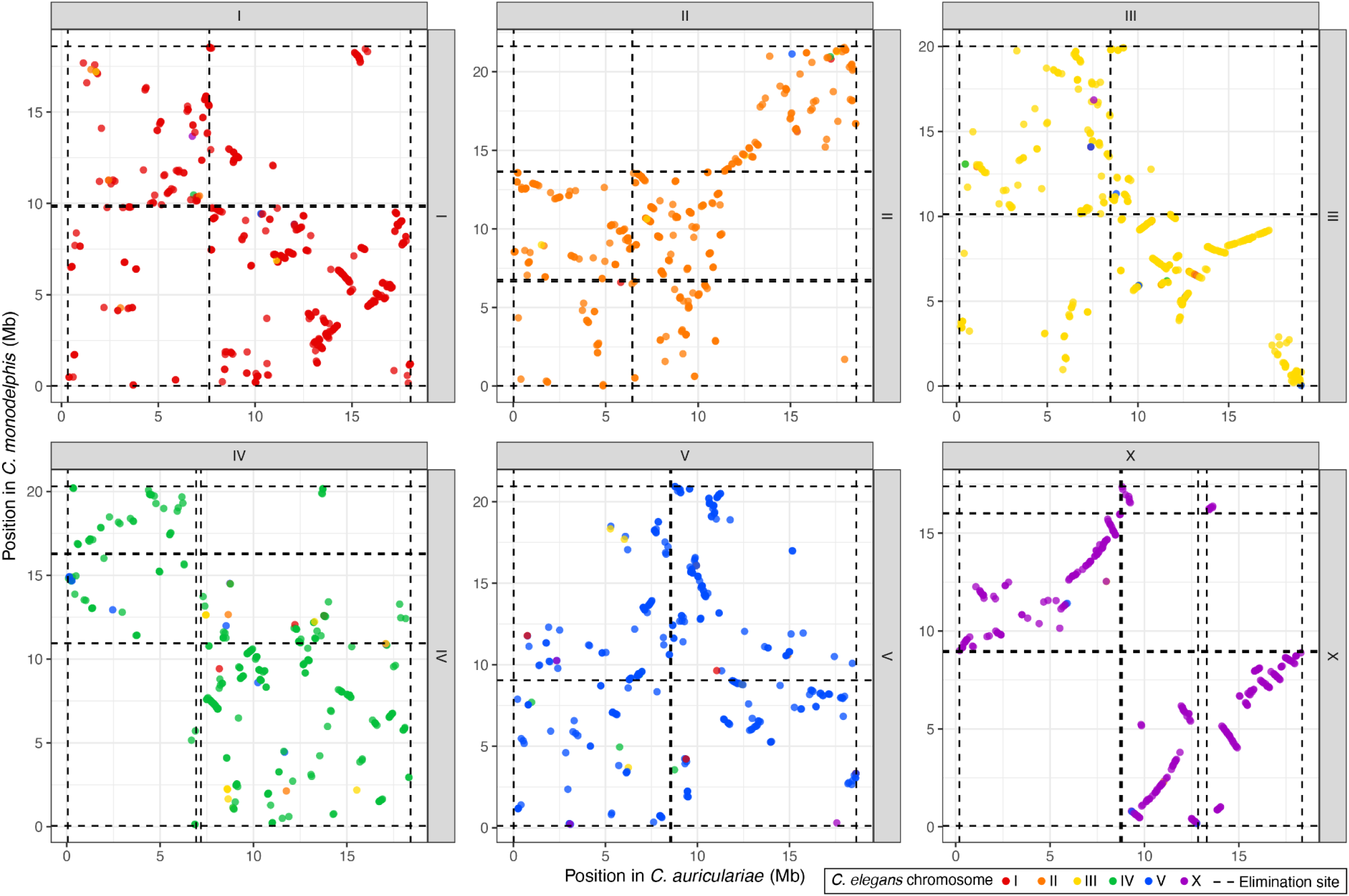
Some elimination sites coincide with sites of rearrangement. Oxford plot showing the relative position of 2,375 BUSCO genes in the six *C. monodelphis* and *C. auriculariae* chromosomes (I-V, X). Genes are coloured by the *C. elegans* chromosome (I-V, X) in which the orthologous gene is found. The locations of telomere addition sites are shown with dotted lines. 64 genes, which were either on different chromosomes in the two species or unassigned to the six *C. elegans* chromosomes, are not shown.

In some autosomes, particularly chromosomes II and IV, we observed domains within which gene order is extensively rearranged, but between which rearrangements are rare (Fig. 5; Fig. S15). Internally eliminated regions sometimes occur at the boundaries of these domains. For example, the genes upstream of an internally eliminated region at ∼13.6 Mb on chromosome II in *C. monodelphis* are highly rearranged in *C. auriculariae* but have almost exclusively remained within the first 12.5 Mb of II, while downstream genes have remained in the last 7.5 Mb (Fig. 5; Fig. S15). Similarly, genes upstream of an internally eliminated region at ∼7.1 Mb on chromosome IV in *C. auriculariae* are highly rearranged in *C. monodelphis* but have remained almost exclusively in the last 9 Mb, whereas the downstream genes are predominantly located in the first 13 Mb (Fig. 5; Fig. S15). We observed similar rearrangement domains between other, similarly diverged species pairs that do not undergo PDE (Fig. S16), suggesting that the domains may form independently of elimination. Gene order was too highly rearranged between *C. parvicauda* and either *C. auriculariae* or *C. monodelphis* (Fig. S14) to assess whether there was any relationship between eliminated regions and rearrangement domains in this species.

To assess whether any eliminated regions or telomere addition sites showed evidence of orthology despite extensive rearrangement, we compared the identities of the three genes located immediately upstream of each telomere addition site. We identified six sites in *C. auriculariae* and *C. monodelphis* that shared at least two genes (one on chromosome I, one on chromosome III, and four on the X chromosome; Table S5). Notably, one of these X-linked sites also shared genes with a site on the *C. parvicauda* X chromosome (Table S5). Specifically, the three genes immediately upstream of telomere addition sites at 8.8 Mb on the *C. auriculariae* X chromosome and 17.4 Mb on the *C. monodelphis* X chromosome were also found immediately upstream of an elimination site at 8.6 Mb on the *C. parvicauda* X chromosome. The genes retained the same relative order across all three species, although they were inverted with respect to the telomere addition site in *C. parvicauda* (Fig. S17). These commonalities, together with the relatively small number of elimination sites in each species, suggest that these telomere addition sites are orthologous.

### PDE is absent in other *Caenorhabditis* species

PDE in *O. tipulae*, which occurs only at the chromosome ends, was detected only through close examination of read coverage in these regions^8^. To determine whether PDE is more widespread within *Caenorhabditis*, we searched the genomes of 30 *Caenorhabditis* species and the closely-related outgroup species *Diploscapter coronatus*, all assembled using long-read data, for evidence of PDE. Specifically, we searched for telomeric repeat-containing long-reads mapping internally to chromosomes, low coverage reads that extend beyond chromosome ends, and/or Hi-C data, where available, indicating more chromosomes than expected (n=6 for *Caenorhabditis*, n=1 for *Diploscapter coronatus*). Except for *C. auriculariae*, *C. monodelphis*, and *C. parvicauda*, the Hi-C contact maps for all other species supported the expected karyotype (Fig. 1; Fig. S1). We detected no sites with telomeric repeat-containing reads mapping internally in 13 of these species, and a further seven species had only one or two such sites (Fig. S18). In *C. japonica* and *C. sinica*, we detected 11 and 9 putative sites, respectively (Fig. S18), but close inspection showed these to be at low coverage or associated with repetitive DNA. Moreover, all were confined to a subset of chromosome ends, despite both species having highly contiguous genome assemblies, and thus were not consistent with PDE.

Taken together, our results suggest that PDE was present in the last common *Caenorhabditis* ancestor and was lost on the branch that separates *C. auriculariae, C. monodelphis*, and *C. parvicauda* from the remaining species.

## Discussion

Here, we report our unexpected discovery of PDE in three early-diverging *Caenorhabditis* species, along with evidence that it was present in the last common *Caenorhabditis* ancestor.

The molecular mechanism of PDE in *Caenorhabditis*, and indeed in any nematode, remains unknown. In *O. tipulae*, chromosome breakage and neo-telomere addition is highly precise and occurs at sites that contain conserved motifs (or SFEs) that are necessary for elimination^9^. Similarly, we found SFEs associated with telomere addition sites in all three *Caenorhabditis* species. The *C. auriculariae* and *C. monodelphis* SFEs consist of two short motifs separated by a variable-length gap and display clear sequence similarity to each other. In contrast, the SFE identified in *C. parvicauda* comprises a single, longer motif with no obvious sequence similarity to those in *C. auriculariae* and *C. monodelphis*. Unlike the *O. tipulae* SFE, the *Caenorhabditis* SFEs are not palindromic and they are typically positioned upstream of the telomere addition sites, rather than spanning the site of telomere addition. We hypothesise that the SFEs are recognised by one or more sequence-specific DNA-binding proteins, which either directly induce double-strand breaks or recruit nucleases to these sites. These same proteins may also facilitate the recruitment of telomerase or associated factors to promote *de novo* telomere addition and stabilise the newly formed chromosome ends.

The biological function of PDE in *Caenorhabditis*, and in nematodes generally, remains unclear. The prevailing hypothesis is that PDE resolves germline-soma conflict by preventing expression of germline-specific genes in somatic tissues^7,38,39^. Consistent with this, *puf-8*, an RNA-binding protein with multiple roles in the *C. elegans* germline^23–25^, is eliminated in both *C. auriculariae* and *C. monodelphis*. Moreover, multiple genes with known germline functions, including post-transcriptional regulation of germline mRNAs, P granule formation, and meiotic pairing, are eliminated in *C. parvicauda.* Several of the eliminated genes in *Caenorhabditis* are responsible for regulating the translation of diverse germline mRNAs, and could therefore affect multiple processes if misexpressed in the soma. However, we see several problems with this explanation of why PDE exists in *Caenorhabditis*. As in *Oscheius* and *Mesorhabditis*^8–10^, only a very small number of genes are eliminated in *Caenorhabditis*, and most are either multi-copy or lack homologs in related species. The vast majority of key germline genes are not eliminated, and control of their expression in the soma is likely achieved by conventional mechanisms involving histone modifications or small RNAs^40–43^. Moreover, we found almost no conservation in the genes that are eliminated in the three species. The germline expression control hypothesis also fails to explain why chromosome ends are consistently removed, despite generally lacking important germline genes, a phenomenon also observed in ascaridids, *Oscheius*, and *Mesorhabditis*^4,8,10^. Future disruption of the as-yet-unidentified elimination machinery might reveal the consequences of retaining the usually-eliminated DNA in the soma, and thus what function PDE serves.

The most parsimonious explanation of the phylogenetic distribution of PDE in *Caenorhabditis* is that it was present in the last common ancestor of the genus and subsequently lost early during the evolution of many species, including *C. elegans*. In support of this, we identified six telomere addition sites with shared upstream genes between *C. auriculariae* and *C. monodelphis*, and one of these sites also shared genes with a site in *C. parvicauda*. Given the relatively small number of telomere addition sites in each genome, the probability of such overlaps arising independently is very low, and these sites are therefore highly likely to be orthologous. Consistent with this, many of the key features of PDE are similar in the three eliminating species. The telomere addition sites are highly precise and are associated with upstream SFEs, and the newly-formed somatic telomere repeat arrays are shorter than those in the germline. DNA is eliminated from both chromosome ends and from within chromosomes, the latter of which means that all three species have distinct germline and somatic karyotypes. The proportion of the genome eliminated in each species is relatively small, and, other than a handful of genes with key germline functions, the majority of eliminated genes are multi-copy or appear to have evolved recently.

However, not all features are conserved between the three species. In particular, the *C. parvicauda* SFE is distinct from the *C. auriculariae* and *C. monodelphis* SFEs. While this could suggest an independent origin in *C. parvicauda*, the deep divergence between *C. auriculariae/C. monodelphis* and *C. parvicauda,* evidenced by complete gene order rearrangement and high protein divergence (average of 1.4 substitutions per site), makes the divergence in SFE structure and sequence less surprising. Another notable difference is in the identities of the eliminated genes across the three species. For example, both *C. auriculariae* and *C. monodelphis* eliminate *puf-8*, whereas *C. parvicauda* eliminates a distinct set of conserved germline genes. However, other than *puf-8*, the overlap in eliminated genes in *C. auriculariae* and *C. monodelphis* is strikingly small, suggesting that the sets of genes eliminated through PDE evolve rapidly. Thus, despite these differences, our results are most consistent with a single origin of PDE in *Caenorhabditis*. Determining whether species at key phylogenetic positions, such as *C. krikudae* (a sister species to *C. auriculariae* and *C. monodelphis*^44^) and *Caenorhabditis* sp. 52 (closer to non-eliminating species than to *C. parvicauda*^45^), also undergo PDE, and whether the elimination machinery is conserved across all three species, will provide important additional evidence for this model.

Since its discovery nearly 140 years ago, PDE has been widely regarded as a biological oddity. Our findings instead suggest that it was a key developmental process in the ancestor of one of biology’s most important model organisms, and one whose mechanisms and functions are only beginning to be understood.

## Methods

### Nematode culture

We obtained *C. monodelphis* JU1667 and *C. auriculariae* NKZ352 from the *Caenorhabditis* Genetics Center (CGC) and *C. parvicauda* NIC534 from Christian Braendle. JU1667 is an inbred derivative of SB341, which was originally isolated from a bracket fungus (*Ganoderma applanatum*) in Berlin, Germany^46,47^. NKZ352 was isolated from a beetle (*Platydema* sp.) in 2015 in Nagoya, Japan^48^. NIC534 is derived from NIC134, which was originally isolated in Sainte-Marie, Madagascar in December 2010^49^. We cultured all three strains on nematode growth media (NGM) plates seeded with *Escherichia coli* strain OP50 and harvested the nematodes by washing the plates with cold M9 into 50 ml Falcon tubes, which we then centrifuged at 4000 rcf for 8 min. We discarded the supernatant and washed the nematodes three times using M9 supplemented with 0.01% Tween, before performing a final wash using PBS buffer. We divided the nematodes into 1.5 ml loBind microtubes (Eppendorf) and recorded the weight of each pellet before flash freezing in liquid nitrogen and storing at −70°C.

### PacBio HiFi sequencing

For *C. auriculariae*, DNA was extracted from a 40 mg frozen worm pellet using a phenol-chloroform protocol with TNES buffer. Worms were ground in a 1.5 ml Eppendorf tube with a pestle in 100 µl TNES buffer, followed by addition of 200 µl TNES and 10 µl Proteinase K (20 mg/ml). The mixture was digested for 2 h at 56 °C and 500 rpm. After digestion, 300 µl phenol/chloroform/isoamyl alcohol (25:24:1) was added, mixed thoroughly with a bore tip, and centrifuged for 10 min at 14,000 rcf and 4 °C. The supernatant was transferred to a new tube, extracted twice with an equal volume of chloroform (mixing by inversion and centrifuging each time under the same conditions). DNA was precipitated with 0.7x isopropanol, incubated for 10 min at room temperature, and pelleted at 14,000 rcf for 15 min at 4 °C. The pellet was washed twice with 70% ethanol and eluted in Qiagen Elution Buffer. High-molecular-weight DNA was sheared twice using a Megaruptor 3 (setting 29; Diagenode), followed by SPRI cleanup and size selection with 0.81× ProNex beads (Promega).

For *C. monodelphis*, several DNA extraction methods were used for PacBio sequencing: the Qiagen MagAttract kit, the Circulomics Nanobind Tissue Big DNA Kit, and two phenol–chloroform extractions (one using TNES buffer and another using CTAB buffer). Using the MagAttract kit from Qiagen, we extracted high molecular weight DNA from a 60 mg frozen pellet of nematodes with the following modifications. The lysis mix (200 µl PBS, 20 µl Proteinase K, 4 µl RNase A, 150 µl Buffer AL) was prepared on ice. We added 75 µl of the lysis buffer mix to the frozen nematode pellet and used a BioMasher II to disrupt the pellet. We added the remaining lysis buffer and mixed with a wide bore tip. We transferred the lysis solution to a 2 ml LoBind microtube (Eppendorf) and digested overnight at 45°C mixing at 600 rpm in a ThermoMixer C (Eppendorf). We added 15 µl of MagAttract Suspension G before mixing with 280 µl Buffer MB. We eluted the MagAttract beads twice using 200 µl of Buffer AE. We incubated the second elution mix at 25°C with 1000 rpm for 3 min in a ThermoMixer C before transferring the elution liquid to a new 1.5 ml LoBind microtube. For the extractions with the Nanobind Tissue Big DNA Kit and the Buffer NL from Circulomics, we used a worm pellet of 70 mg. We followed the provided “Nanobind High Molecular Weight C. elegans DNA Extraction Protocol” with the following modifications. We added 20 µl Proteinase K and 150 µl Buffer NL to the frozen pellet and disrupted the tissue with the BioMasher II. We eluted the DNA from the nanobind disk overnight in 200 µl EB buffer (Qiagen). We then mixed the extracted DNA several times with a bore pipette tip before proceeding to QC measurements. For the CTAB phenol-chloroform extraction we used a frozen worm pellet of 50 mg. We added 1 µl of beta-mercaptoethanol to 1 ml CTAB buffer. We then added 100 µl of CTAB buffer and used a pestle to grind the frozen tissue. After that we added 400 µl more of the CTAB buffer. We added 20 µl Proteinase K (20 mg/ml) and digested the tissue for 1 hr at 56 °C and 500 rpm in a ThermoMixer C (Eppendorf). We added 10 µl RNaseA, mixed well with a bore tip, and let it incubate for 10 min at room temperature. We then added 500 µl of a phenol/ chloroform/isoamyl alcohol mix (25:24:1). We centrifuged the mix for 10 min at 14,000 rcf at 4 °C. We transferred the supernatant to a new tube and added 1x chloroform. We mixed well by inversion. We centrifuged the tube for 10 min at 14,000 rcf at 4 °C. The supernatant was taken to a new tube and the chloroform extraction was repeated. We then added 0.7x isopropanol to the supernatant. The tube was inverted several times and left for 10 min at room temperature. We pelleted the DNA at 14,000 rcf for 15 min at 4 °C before performing two ethanol washes with 70% ethanol. The DNA pellet was eluted in EB.

For *C. parvicauda*, we extracted DNA from a 20 mg frozen worm pellet using the Monarch® HMW DNA Extraction Kit for Tissue (NEB) with the following modifications. We added 50 µl of the lysis buffer mix to the frozen worm pellet and did the pestle step. We then added the rest of the lysis buffer and transferred the mix in a 2 ml DNA LoBind Tube (Eppendorf). The mix was digested at 56 °C in a ThermoMixer C (Eppendorf) at 600 rpm for 2 hr. After RNA digestion we spun the sample for 5 min at 2500 rcf and transferred the supernatant to a new 1.5 ml DNA LoBind Tube (Eppendorf) leaving about 30 µl behind. We added 30 µl of lysis buffer to the mix and put the sample for 5 min on ice. We added 300 µl cold protein separation solution and inverted the sample at about 20 rpm for 1 to 2 min. We put the sample for 10 min on ice. We spun the sample at 16,000 rcf for 25 min at room temperature. We transferred the supernatant to a new 2 ml DNA LoBind Tube (Eppendorf) and added 3 glass beads. We added 550 µl isopropanol and inverted the sample several times before we put it in a wheel for 5 min at 7 rpm. From here we followed the recommended protocol steps. We sheared the extracted high molecular weight DNA with the Megaruptor 3 twice (setting 28) (Diagenode). The sheared DNA was SPRI cleaned and size-selected with 0.81x of ProNex beads (Promega).

PacBio libraries were prepared from extracted DNA by the Scientific Operations: Sequencing Operations core at the Wellcome Sanger Institute using the PacBio Low DNA Input Library Preparation Using SMRTbell Express Template Prep Kit 2.0. Libraries were sequenced on PacBio Single Sequel IIe flow cells.

### Oxford nanopore sequencing

*C. auriculariae* genomic DNA was extracted using Genomic-tip (Qiagen) following the manufacturer’s protocol. Two micrograms of genomic DNA was used to prepare Nanopore sequencing library using the Ligation Sequencing Kit and sequenced with a single 24 h run with a MinION flowcell according to the manufacturer’s protocol (Oxford Nanopore Technologies). Base calling for the runs was performed with Guppy v.3.1.5 using the ‘dna_r9.4.1_450bps_fast’ model.

### Hi-C sequencing

Hi-C library preparation and sequencing was performed by the Scientific Operations Sequencing Operations core at the Wellcome Sanger Institute for all three species. Pellets of mixed-stage nematodes weighing approximately 20-60 mg were processed using the Arima Hi-C version 2 kit following the manufacturer’s instructions. Illumina libraries were prepared using the NEBNext Ultra II DNA Library Prep Kit and sequenced on one-eighth of a NovaSeq S4 lane using paired-end 150 bp sequencing.

An additional Hi-C library for *C. auriculariae* was constructed from about 1000 mixed-stage nematodes using an Arima Hi-C+ kit (Arima Genomics Inc.) and a Collibri ES DNA library prep kit (Thermo Fisher Scientific) according to the manufacturers’ protocols and was sequenced using a MiSeq instrument with the MiSeq reagent kit v3 (101 cycles × 2).

### Assembly and curation of the *C. auriculariae* genome

We assembled ONT reads using NextDenovo version 2.4.0 (https://github.com/Nextomics/NextDenovo), followed by three rounds of base error correction using Pilon version 1.24^50^ with Illumina short reads. We used the Hi-C data to scaffold the assembly with Juicer (version 1.5)^51^ and 3D-DNA pipeline v180114^52^ with parameters -g 500 -e. We used Juicebox v1.11.08^53^ for manual curation of the assembly. We manually extended the *C. auriculariae* chromosome ends by aligning PacBio HiFi reads to the scaffolded assembly using minimap2 v2.26^54^. We then used Geneious Prime (https://www.geneious.com) to identify reads or contigs that extended beyond the existing scaffold ends.

### Assembly and curation of the *C. monodelphis* genome

We removed adapter sequences from the PacBio HiFi data using HiFiAdapterFilt^55^. We used Jellyfish 2.3.0 ^56^ to count k-mers of length 31 in each read set and GenomeScope 2.0^57^ to estimate genome size and heterozygosity. To prevent assembly fragmentation at sites of telomere addition, we identified and removed reads containing at least 10 consecutive occurrences of the nematode telomeric repeat sequence (TTAGGC) using seqkit v2.1.0^58^. We assembled the remaining reads using hifiasm v0.16.1-r375^59^ (--primary). We randomly subsampled 10% of the Hi-C reads using SAMtools 1.14^60^ and aligned them to the hifiasm primary assembly using bwa-mem 0.7.17-r1188^61^, filtered out PCR duplicates using picard 2.27.1-0 (available at http://broadinstitute.github.io/picard/), and scaffolded the assembly using YaHS^62^. We ran BlobToolKit 2.6.5^63^ on the scaffolded assembly and used the interactive web viewer to manually screen for scaffolds derived from non-target organisms. We removed 131 scaffolds that corresponded to Proteobacteria, Actinobacteria, or Uroviricota from the assembly. We then used MitoHiFi 2.2^64^ to identify and annotate the mitochondrial genome. We removed residual haplotypic duplication from each assembly using purge_dups 1.2.5^65^. However, to avoid removing contigs that corresponded to eliminated DNA, we only removed contigs labelled as haplotigs and retained those labelled as junk or repeats. We then scaffolded the purged primary assembly using Hi-C data and YaHS (using the --no-contig-ec parameter to prevent contigs being broken), as previously described.

Although removing the telomere-containing reads from hifiasm assembly prevented fragmentation at sites of telomere addition, it meant that the scaffolds did not end in telomeric repeat arrays. We used a variety of different approaches and evidence to manually extend the *C. monodelphis* chromosome ends. We generated hifiasm assemblies using all reads and only reads containing telomeric repeat sequences, which we then aligned to the scaffold assembly using minimap2 2.24-r1122^54^ (-cx asm5 --secondary=no --cs). We also aligned PacBio HiFi reads to the scaffolded assembly using minimap2 (-ax map-hifi). We then used IGV to visualise read and assembly alignments and identify reads or contigs that extended beyond the existing scaffold ends. Where existing scaffolds terminated in unique sequence, we used CAP3 v02/10/15^66^ to directly join aligned reads or contigs to the scaffold ends. Where existing scaffolds terminated in repetitive sequence, we used agptools^67^ to add these reads or contigs to the scaffold with a gap containing 200 N characters. Misassembled sequence was removed using the getfasta function in BEDtools v2.30.0^68^. All assembly corrections were assessed using Dotter^69^ and by inspecting read alignments using IGV 2.18.4^70,71^. A detailed list of changes made is available in Supplementary Text 1.

The final assembly was checked for residual contamination and corrected using BlobToolKit^63^, and FCS^72^. Manual curation was performed using JBrowse 2^73^, HiGlass^74^ and PretextView (available at https://github.com/wtsi-hpag/PretextView), where we made three breaks, thirteen joins, and removed six haplotigs. Final metrics were calculated using seqkit and BUSCO 5.2.2^20^ (with ‘metaeuk’ as the gene predictor and ‘nematoda_odb10’ as the lineage).

### Assembly and curation of the *C. parvicauda* genome

We removed adapter sequences from the PacBio HiFi data using HiFiAdapterFilt. We used Jellyfish 2.3.0 to count k-mers of length 31 in each read set and GenomeScope 2.0 to estimate genome size and heterozygosity, which revealed that the strain we sequenced was highly polymorphic (Fig. S2). We initially assembled the genome using hifiasm v0.16.1-r375, but the resulting assembly had low BUSCO completeness (72.4%; metaeuk and nematoda_odb10) and was highly fragmented (contig N50 of 382 kb). We reassembled the data using three other assemblers Flye 2.9-b1768^75^ (with the parameters --pacbio-raw and -g 100m), wtdbg2 0.0^76^ (with the parameters -x sq), and metaMDBG 0.3^77^ (with default parameters). Although all assemblies were highly fragmented (N50s of 149 kb - 290 kb), the metaMDBG assembly had the highest BUSCO completeness (77.7%, compared with 74.3 for Flye and 71.8% for wtdbg2), and so we selected this as the primary assembly. We used Hi-C data to scaffold the assembly, as previously described, and ran BlobToolKit 2.6.5 to screen for scaffolds derived from non-target organisms. We removed 175 contigs that were labelled as Proteobacteria or Actinobacteria. We then used MitoHiFi 2.2 to extract the mitochondrial genome and purge_dups 1.2.5 to remove haplotigs, as previously described. We then scaffolded the purged primary assembly using Hi-C data and YaHS (using the parameter that prevents contig breaking), as previously described. Residual contamination checks and manual curation were performed as previously described, where we made 186 breaks, 411 joins, and removed 49 haplotigs.

### Protein-coding gene prediction

We predicted protein-coding genes in 22 *Caenorhabditis* species and *Diploscapter coronatus* (Table S6). We supplied RNA-seq datasets for each species (Table S6), along with OrthoDB v11 nematoda dataset^78^ as protein homology evidence, to BRAKER3 with the following modifications/parameters: (1) we modified the TSEBRA step within BRAKER3 to remove single-exon genes (--filter_single_exon_genes) and ignore phase when grouping transcripts (--ignore_tx_phase); (2) we supplied a custom TSEBRA config file (available at https://github.com/lstevens17/caenoPDE/); (3) we allowed the prediction of alternative transcripts when supported by evidence (--alternatives-from-evidence=true); (4) we allowed non-canonical splice sites when supported by evidence (--allow_hinted_splicesites=gcag,atac); (5) we allowed overlapping genes on different strands (--singlestrand=true); (6) we required all gene models to be complete genes (--genemodel=complete); (7) we required a minimum intron length of 15 bp (--min_intron_len=15). Finally, we assessed the completeness of each gene set using BUSCO 5.2.2^20^ (using the nematoda_odb10 lineage and -mode proteins) (Table S6).

### *Caenorhabditis* phylogeny

To infer the *Caenorhabditis* phylogeny, we collated genomes for 30 *Caenorhabditis* species and *Diploscapter coronatus* (all of which were chromosome-level, apart from *C. bovis*)^12,14–18,79–82^. We ran BUSCO 5.2.2 (using the nematoda_odb10 lineage) on all genomes and extracted the protein sequences of 1,601 BUSCO genes that were present and single copy in at least 30 of the 31 species using busco2fasta (available at https://github.com/lstevens17/busco2fasta). We aligned the sequences using MAFFT v7.526^83^ and inferred gene trees for each alignment under the LG substitution model^84^ with gamma distributed rate variation among sites (LG+G4) using IQ-TREE 2.2.0.3^85^. We provided the resulting gene trees to Astral v5.7.8133^86^ to infer the species tree. To estimate branch lengths in amino acid substitutions per site, we concatenated the alignments of each single-copy orthogroup into a supermatrix using catfasta2phyml v1.1.0 (available at https://github.com/nylander/catfasta2phyml) and used IQ-TREE to estimate branch lengths under the general time reversible substitution model with gamma-distributed rate variation among sites. We visualised and rooted the resulting species tree using the iTOL webserver^87^.

### Identification of telomere addition sites and eliminated regions

To identify telomere addition sites, we aligned PacBio HiFi reads to their corresponding reference genomes using minimap2 2.24-r1122^54^ (-ax map-hifi). The resulting BAM files were supplied to delfies 0.9.0^88^ (--min_supporting_reads 1) to identify sites where mapped reads contained telomeric repeat sequence that was soft-clipped. These candidate sites were then manually inspected using IGV 2.18.4^70,71^ to confirm the presence of elimination. In a subset of *C. auriculariae* and *C. monodelphis* sites, the eliminated DNA had sequence similarity to the telomeric repeat, and the exact telomere addition site was ambiguous. In these cases, we defined the first eliminated base as the base immediately downstream of the first GG dinucleotide encountered after the telomere-homologous region, consistent with patterns observed at other sites.

### Identification of motifs at telomere addition sites

We used BEDtools to extract regions surrounding each telomere addition site, accounting for strand orientation. In *C. auriculariae* and *C. monodelphis*, both of which have SFEs composed of two motifs separated by a gap, we extracted two regions upstream of each telomere addition site. To identify motif-1, we extracted sequences spanning bases 20 to 70 bp upstream of each site (a 50 bp region of retained DNA) in *C. auriculariae* and 20 to 100 bp upstream (an 80 bp region) in *C. monodelphis*. To identify motif-2, we extracted 30 bp of upstream retained DNA and 10 bp of downstream eliminated DNA (a 40 bp region) in both species. We excluded the left end of chromosome V in *C. auriculariae* (where the assembly is incomplete as it contains the ribosomal DNA repeat array) and the two variable sites in *C. monodelphis.* For *C. parvicauda*, which had a single motif, we extracted 100 bp of upstream retained DNA and 10 bp of downstream eliminated DNA (a 110 bp region). We then identified overrepresented sequences in all three species using the MEME suite v5.5.7^21^, limiting the search to the positive strand only. To identify motif-1 in *C. auriculariae* and *C. monodelphis*, we searched for a motif that was between 11 and 16 bp in length that occurred in all sites. To identify motif-2, we searched for a motif that was 11 bp in length that occurred in all sites. For *C. parvicauda*, we searched for one motif between 10 and 30 bp in length that occurred in all sites. To determine the specificity of the identified motifs, we searched for instances of each motif in the corresponding genomes using FIMO v5.5.7^89^. For *C. auriculariae* and *C. monodelphis*, we used a custom Python script (find_motif_pairs.py) to identify pairs of motifs that met the following criteria: (i) both motifs were located on the same chromosome; (ii) the motifs did not overlap; (iii) when motif-2 occurred on the positive strand, motif-1 was located upstream of motif-2, and when motif-2 occurred on the negative strand, motif-1 was located downstream. We used a second custom Python script (filter_duplicate_memes.py) to discard motif pairs in which either occurrence overlapped with a higher-scoring pair. For *C. parvicauda*, we used the FIMO output directly. We classified each motif occurrence as being ‘real’ (associated with a telomere addition site), ‘retained’ (occurring within retained DNA), and ‘eliminated’ (occurring within eliminated DNA) using BEDtools. Motifs that had scores lower than the lowest-scoring real SFE in each species and/or motifs that had gap sizes greater than 50 bp (*C. auriculariae*) and 70 bp (*C. monodelphis*) were filtered out. We also filtered out any motifs that occurred within germline telomere repeat blocks.

### Comparison of germline and somatic telomere lengths

To identify germline telomere-repeat containing reads, we used BEDtools intersect to extract reads mapped to the first and last 100 bp of each germline chromosome end. We then used the ‘locate’ function in seqkit to find reads containing at least ten consecutive copies of the nematode telomeric repeat. To identify somatic telomere-repeat containing reads, we applied the same procedure but extracted reads located 100 bp upstream of each telomere addition site. Germline telomere-containing reads could not be identified in *C. parvicauda* because the germline chromosome ends were not resolved in our assembly.

### Analysis of eliminated genes

To identify eliminated genes, we used BEDtools intersect (-f 1) to identify genes that were contained within the previously identified eliminated regions. To infer the relationship of these genes to other genes in the genome or in other *Caenorhabditis* species, we used an orthology clustering approach. We first collated protein sequence and GFF3 files for 53 *Caenorhabditis* and two *Diploscapter* outgroup species and used AGAT v0.8.0^90^ (agat_sp_keep_longest_isoform.pl) to identify the longest isoform of each protein-coding gene. We then clustered the isoform-filtered proteomes using OrthoFinder v2.5.4^91^ (-og). We also used InterProScan 5.61-93.0^92^ (-dp -t p --appl Pfam --goterms) to identify Pfam domains in proteins encoded in eliminated genes. We used a custom script (summarise_orthogroups.py) to summarise the relationship of each eliminated gene to other genes in the genome and other species, the output of which can be found in Data S2.

### Identification and enrichment of satellite repeats

To identify satellite repeats in the genomes of *C. auriculariae*, *C. monodelphis,* and *C. parvicauda*, we ran Tandem Repeats Finder (TRF) 4.09^93^ with the following parameters: match = 2, mismatch = 7, delta = 7, PM = 80, PI = 10, minscore = 50, maxperiod = 500. We filtered the TRF output to retain repeat units that were ≥50 bp in length, occurred ≥100 times in the genome, and had an average nucleotide identity of ≥85%. To reduce redundancy, we clustered the filtered repeats using VSEARCH v2.22.1^94^ with a sequence identity threshold of 90% and kept one representative per cluster (the centroid). The centroid sequences were then searched against their respective genome with BLASTN 2.5.0+^95^, with a maximum e-value of 0.01. We then provided these BLAST hits to BEDtools v2.30.0^68^ to calculate the coverage of each satellite repeat unit in 50 kb windows.

To determine if satellite repeats were enriched in the eliminated regions, we used BEDtools to merge adjacent satellite repeat locations and summed the length of the merged intervals to calculate total genome-wide satellite repeat content. We used BEDtools coverage to calculate the amount of satellite DNA that is eliminated. To calculate fold enrichment, we compared the observed span of eliminated satellite repeat to the expected span (calculated by multiplying the genome-wide satellite repeat proportion by the total eliminated span) (Table S4). To determine if satellite repeats were statistically significantly enriched, we applied the overlapPermTest function from the regioneR v1.38.0 R package^96^, using 1,000 permutations. Each permutation randomly locates the eliminated regions across the genome and calculates their overlap with satellite repeats. The overlap between satellite repeats and the eliminated regions was compared against the distribution of permuted overlaps to generate z-scores and empirical p-values.

### Identification and enrichment of transposable elements

We used Earl Grey v6.1.1^97^ (-r rhabditida -e) to identify transposable elements (TEs) in the genomes of three *Caenorhabditis* species that undergo PDE, as well as in 26 other *Caenorhabditis* species with chromosome-level reference genomes. To assess TE enrichment within eliminated regions, we used regioneR as previously described. Enrichment analyses were performed both globally (using all TEs) and separately for each TE superfamily.

To test whether Helitron-superfamily transposable elements (TEs) were significantly enriched at chromosome ends in species that do not undergo genome elimination, we first calculated the average proportion of each chromosome that is eliminated from both ends in the three species that undergo elimination (2.69% per end). We used a custom Python script (first_and_last.py) to calculate the coordinates of the first and last 2.69% of each chromosome for all species, These regions, along with the locations of all elements belonging to the RC/Helitron superfamily, were then used as input to regioneR to perform permutation tests and assess enrichment, as previously described.

### DNA FISH

FISH was performed with modifications based on a previous protocol for *C. elegans* embryos^98^. Adult female *C. auriculariae* were placed in M9 buffer in a watch glass and incubated for 2-3 hours to obtain embryos at different developmental stages. Worms were then dissected in 10 µl of 1 mM levamisole on poly-L-lysine-coated slides, freeze-cracked, and fixed in pre-chilled methanol (−30°C) in a fixation jar for 10 min. Each slide was washed twice in PBST (0.05% Tween-20 in 1X PBS buffer), followed by incubation with 40 µl of fixative solution (3.7% formaldehyde, 0.8% methanol in 1X PBS) for 10 min. After a PBST wash, slides were treated with 2X SSCT (2X saline-sodium citrate [SSC] buffer [Thermo Fisher Scientific: 15557–04] and 0.5% Triton X-100). They were incubated in denaturation buffer (50% formamide in 1X SSCT) at 60°C for 30 min, then overlaid with 10 µl of probe mixture (10% dextran sulfate, 2X SSC, 50% formamide, 1 μM of DNA probe [Ca_SR2: 5’-Alexa488-GTGGACAAAAATCGACAAAACAAAGACC-3’ (Eurofins), Ca_SR8: 5’-CY3-CAGTTTTGACGTCATCAGTTCAA-3’ (Eurofins), Telomere: 5’-CY3-TTAGGCTTAGGCTTAGGCTTAGGC-CY3-3’, (Integrated DNA Technologies)], 100 μg/ml of RNase A and 0.1% Tween-20). The slides were then covered with coverslips and sealed with Fixogum (Marabu). Samples were denatured at 93°C for 2 min, and hybridized for 3 h at 37°C. The slides were washed twice in 2X SSCT at 37°C and once in 2X SSC at room temperature, then mounted in ProLong Gold with DAPI (Thermo Fisher: P36941).

### Construction of a transgenic strain

The transgenic *C. auriculariae* strain (SA1864, *tjIs391[Cau-eft-3p::GFP::Cau-Histone H2B, HygR]*) that expresses GFP::Histone H2B under the control of the ubiquitous *Cau-eft-3* promoter was established by microparticle bombardment. The plasmid pNH73 for microparticle bombardment was constructed as follows. DNA fragments corresponding to the putative promoter region (686 bp) of the *C. auriculariae eft-3* gene (Cau-eft-3; CAUJ_0077810), GFP, and Cau-Histone H2B including its 3’ UTR region (CAUJ_01174300.t1; [1] ORF + 898 bp), both of which were obtained from a previous version of the *C. auriculariae* reference genome^48^, were PCR-amplified and inserted into the multicloning site of the plasmid pCFJ1662 (Addgene #51482, deposited by Erik Jorgensen), which contains the hygromycin B resistance gene cassette^99^.

Microparticle bombardment of *C. auriculariae* was performed following an established protocol for *C. elegans* and related nematodes^100–102^, with minor modifications. Briefly, L4 and adult worms cultured at 20°C were collected in M9, and subjected to five bombardment shots (approximately 10,000 worms/shot). Two days later, hygromycin B (final concentration 0.33 mg/mL; Invitrogen) was added, and the worms were incubated at 20°C. Surviving worms exhibiting GFP fluorescence were isolated and crossed with wild-type worms for maintenance.

### Microscopy

Fluorescence images were acquired using an Orca-R2 digital CCD camera (Hamamatsu Photonics) mounted on an IX71 microscope (Olympus) equipped with a CSU-X1 spinning disc confocal system (Yokogawa Electric Corporation). A UPlanSApo 100X/1.40 oil-immersion objective lens (Olympus) was used for FISH, and a UPlanSApo 60X/1.30 silicone-immersion objective lens (Olympus) for live imaging. MetaMorph software (version 7.10.3.279, Molecular Devices) was used to control the microscope. Images were collected as Z-stacks at 0.5 µm steps for FISH and 1 µm steps for live imaging at 1 min intervals, and processed with ImageJ software (NIH).

### Genome rearrangement

To compare gene order between a given pair of species, we ran BUSCO on both genomes as previously described. We then used a custom Python script (ce_oxford.py) to retrieve the positions of BUSCO genes that were single copy and complete in each species, along with which *C. elegans* chromosome the gene is found in. For the comparison between *C. auriculariae* and *C. monodelphis*, we used a custom Python script (rename_by_scu.py) to identify the somatic chromosome (e.g. I_1) that each gene was found in. To determine whether rearrangement domains were observed in species that did not undergo PDE, we used the *Caenorhabditis* phylogeny to find pairs of species that showed similar divergence to *C. auriculariae* and *C. monodelphis* (average of 0.883 amino acid substitutions per site). The pair that were most similar were *C. sulstoni* and *C. uteleia* (average of 0.882 substitutions per site).

### Identification of orthologous telomere addition sites

To identify potentially orthologous telomere addition sites, we compared the identities of the three genes located immediately upstream of each site. To do this, we converted the BUSCO full_table.tsv files into BED files and then used a custom Python script (find_upstream_genes.py) to extract the coordinates, orientations, and names of the three genes upstream of each elimination site. We used a second Python script (compare_upstream_overlap.py) to identify sites that shared genes across a given pair of species. Sites with two or more overlapping genes are listed in Table S5. One site on the X chromosome was shared across all three species (Fig. S17). For this site, the coordinates and orientation of 1541at6231 in *C. monodelphis* were obtained from a second BUSCO run using Augustus as the gene predictor (specified with the --augustus parameter) because the gene was labeled “Missing” in the original BUSCO run, which used MetaEuk as the gene predictor.

### Search for PDE across *Caenorhabditis*

We collated chromosome-level reference genomes for 29 species of *Caenorhabditis,* along with genomes for *C. bovis* (which is near chromosome-level) and a chromosome-level genome for the outgroup *Diploscapter coronatus*. All species had either long-read PacBio CLR, PacBio HiFi, or ONT data available. We aligned the long-reads to the corresponding reference genome using minimap2 (specifying map-pb, map-hifi, or map-ont depending on the available data). The resulting BAM files were supplied to delfies to identify sites where mapped reads contained telomeric repeat that was soft-clipped (labelled as S2G) or mapped reads that extend beyond the end of a contig/chromosome containing telomeric repeat sequence (G2S). To exclude S2G sites that were supported by a minority of reads, we used mosdepth 0.3.3^103^ to calculate the median read coverage in each species, and filtered out any sites that had less than 25% of the median coverage. Any S2G sites that passed these filters, in addition to any identified G2S sites, were manually inspected in IGV.

## Data availability

The *C. auriculariae*, *C. monodelphis*, and *C. parvicauda* reference genomes have been deposited in ENA under the BioProject accessions PRJEB40642, PRJEB77269 and PRJEB73705, respectively. PacBio and Hi-C data have been deposited in ENA under the BioProject accessions PRJEB40642 and PRJEB36817. Accession numbers for the genome assemblies and annotations used are available in Table S1 and Table S6. Code, data files, and gene prediction files can be found in the GitHub (https://github.com/lstevens17/caenoPDE).

## Acknowledgements

This work was supported by Wellcome Trust grant 220540/Z/20/A, Japan Science and Technology Agency (JST), Core Research for Evolutionary Science and Technology (CREST) Grant Number JPMJCR18S7 and JPMJCR24B4, and Japan Society for the Promotion of Science (JSPS) KAKENHI grants JP23K17400 and JP24H00549. For the purpose of Open Access, the authors have applied a CC BY public copyright licence to any Author Accepted Manuscript version arising from this submission. We thank the Sanger Scientific Operations Long Read team for sequencing and the Genome Reference Informatics Team in Tree of Life for genome curation assistance. We also thank other members of the Blaxter laboratory, the Sugimoto laboratory, the Kikuchi laboratory, Marie Delattre and Brice Letcher for valuable feedback on the manuscript and analyses. We thank the *Caenorhabditis* Genetics Center, which is funded by NIH Office of Research Infrastructure Programs (P40 OD010440), and Christian Braendle for providing nematode strains.

