## Supplementary Material for "Programmed DNA elimination was present in the last common ancestor of *Caenorhabditis* nematodes"

The PDF file includes:

Figs. S1 to S18
Tables S1 to S6
Supplementary Text

Other Supplementary Materials for this manuscript include the following:

Data S1 to S2
Movies S1 to S2

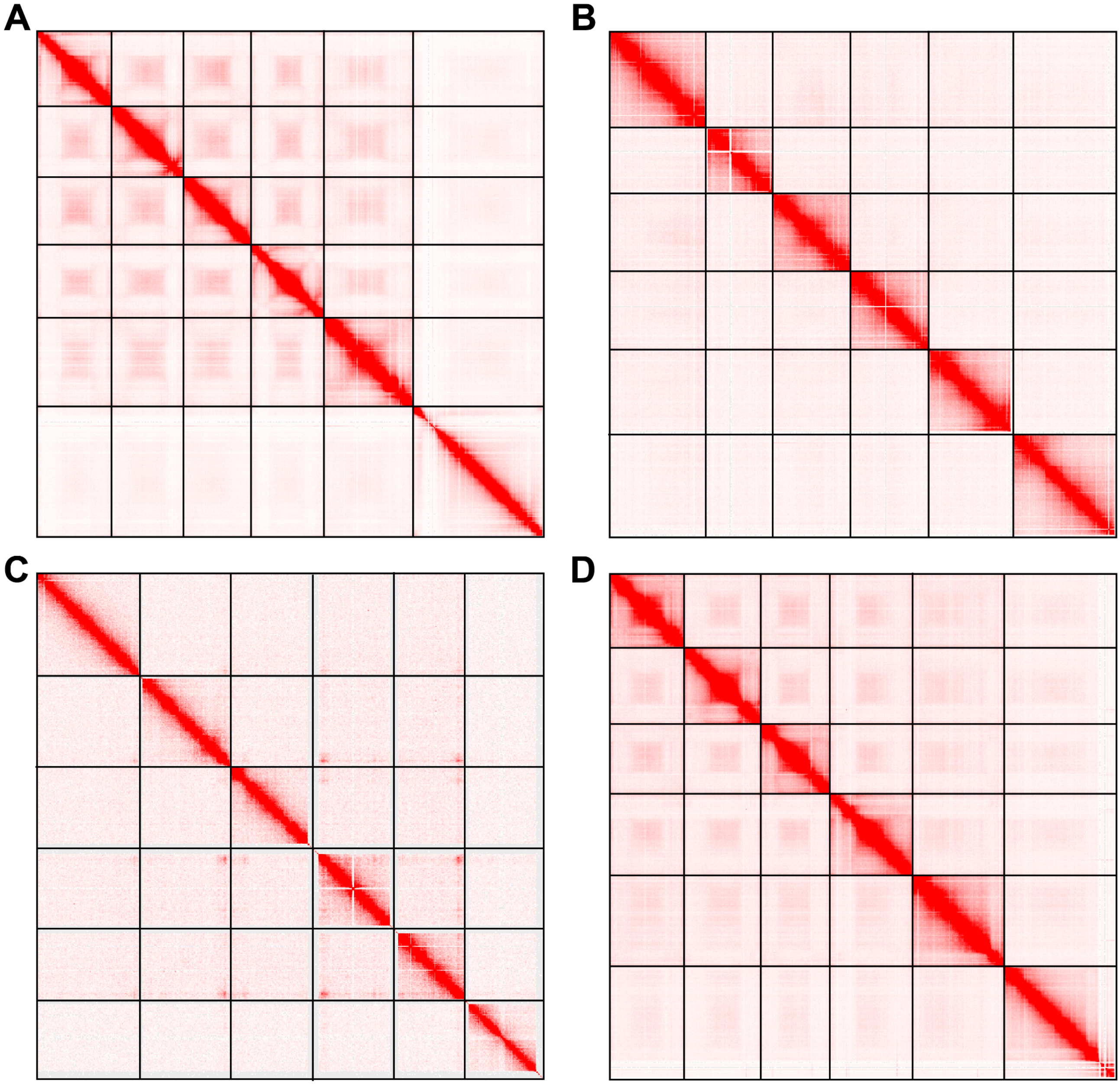


**Fig. S1: Hi-C contact maps for four *Caenorhabditis* species**

Hi-C contact maps for (A) *C. afra*, (B) *C. drosophilae*, (C) *C. plicata*, and (D) *C. wallacei*. Maps were visualised with Juicebox. Chromosomes are ordered I-V, X.

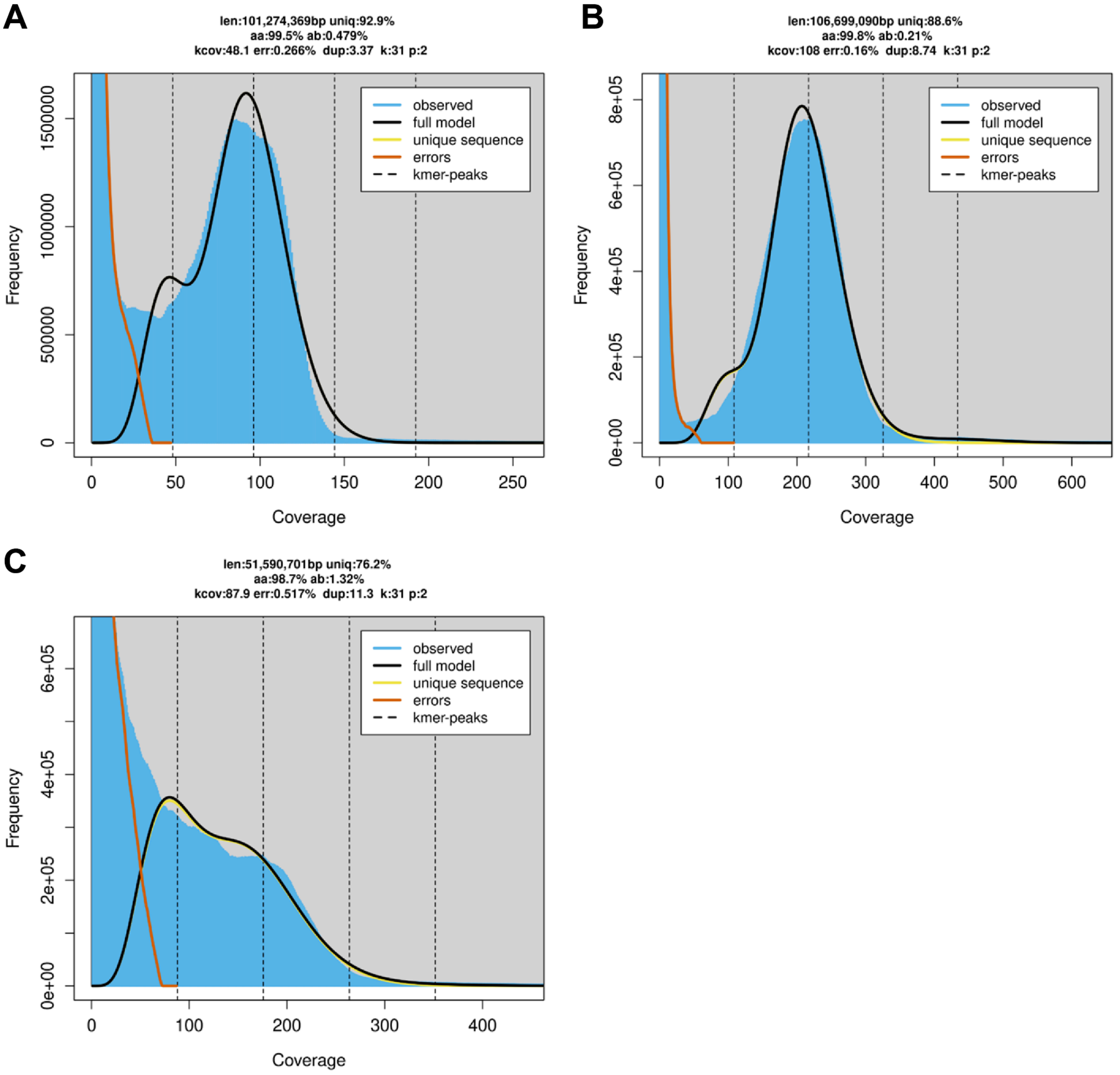


**Fig. S2: K-mer spectra of *C. auriculariae, C. monodelphis*, and *C. parvicauda***

K-mer spectra (*k* = 31) and fitted models for PacBio HiFi reads from GenomeScope 2.0 for (A) *C. auriculariae*, (B) *C. monodelphis*, and (C) *C. parvicauda*. K-mers were counted after removing reads derived from contaminants.

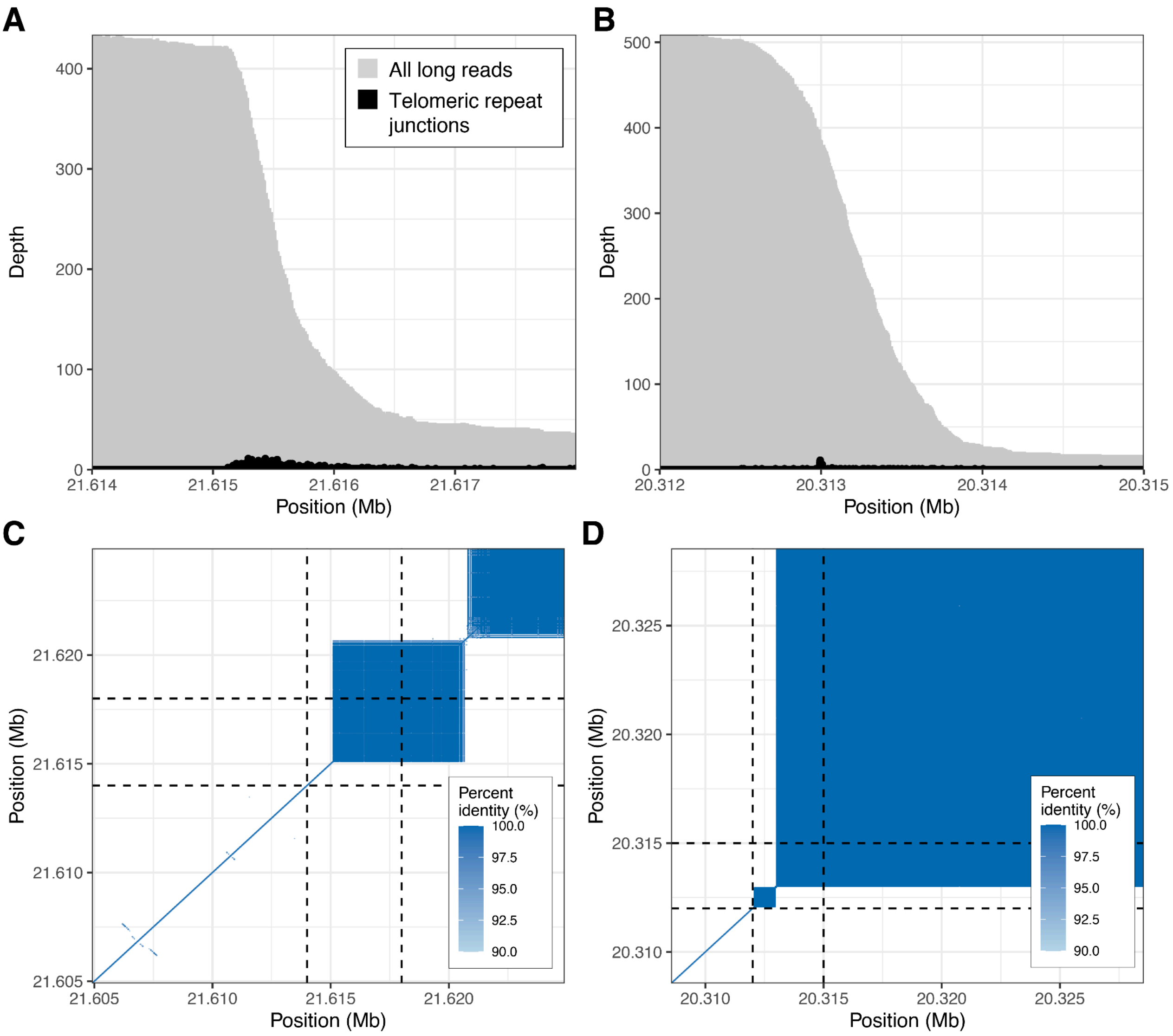


**Fig. S3: Variable telomere addition sites in *C. monodelphis***

Variable telomere addition sites at the right end of chromosome II (A) and IV (B) in *C. monodelphis*. Depth of all PacBio HiFi reads and of junctions between unique sequence and soft-clipped telomeric repeat are shown. Dot plots of the last 20 kb of chromosome II (C) and IV (D) showing satellite repeat arrays occurring immediately upstream of the telomeric repeat arrays. The regions shown in (A) and (B) are indicated with dotted lines. We found multiple low-scoring Cm-SFE occurrences in the repeat arrays (scores of 1.1 in chromosome II and 18.8 in chromosome IV) that potentially explain the variable telomeric repeat addition.

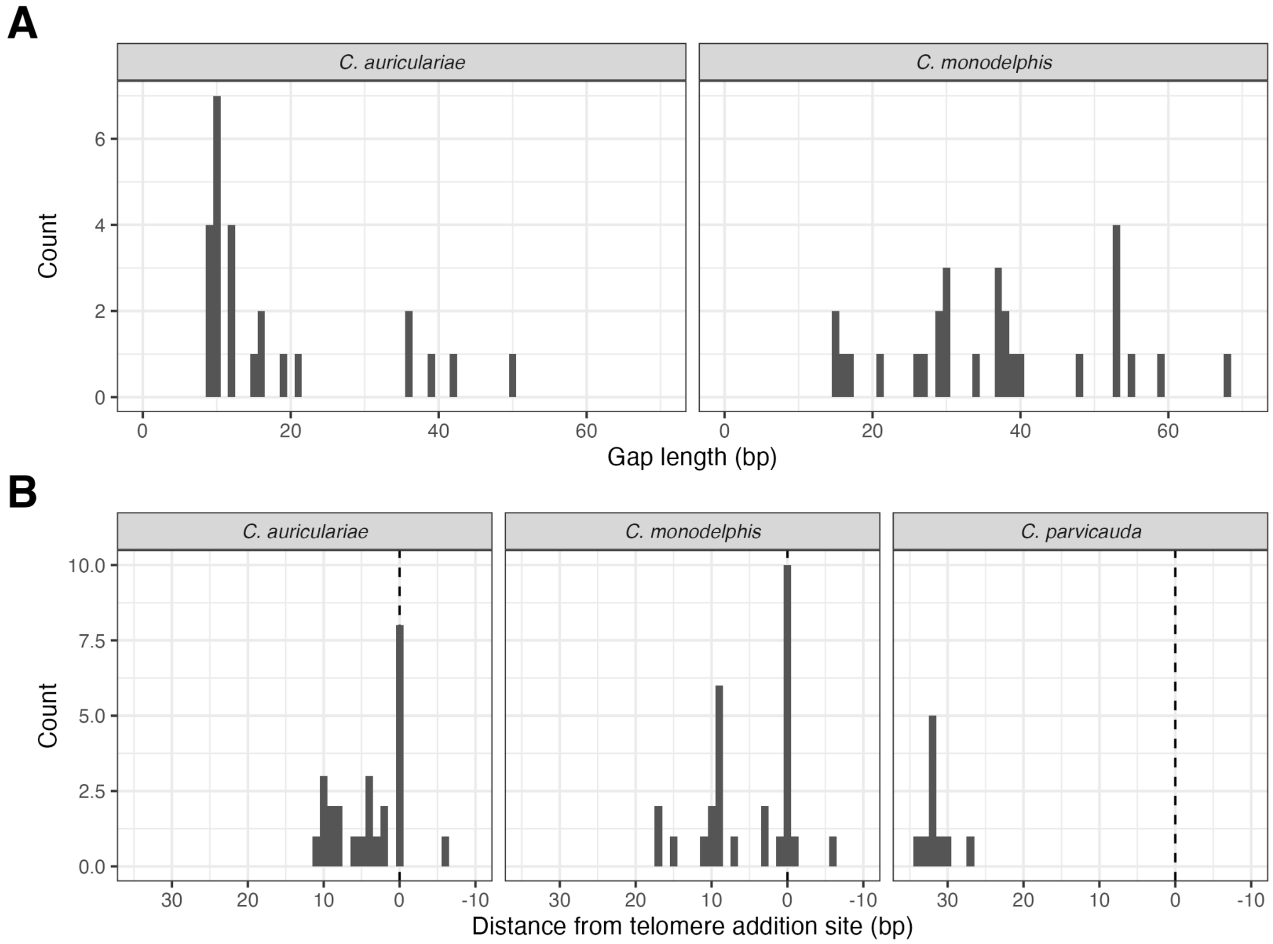


**Fig. S4: Sequence for elimination (SFE) gap size and distance from break site**
(A) Distribution of gap sizes in the *C. auriculariae* and *C. monodelphis* SFEs. (B) Distance between the last base pair of SFEs and the break site. Positive values indicate that the last base pair of the SFE is upstream of the break site; negative values indicate that the last base pair of the SFE is downstream of the break site.

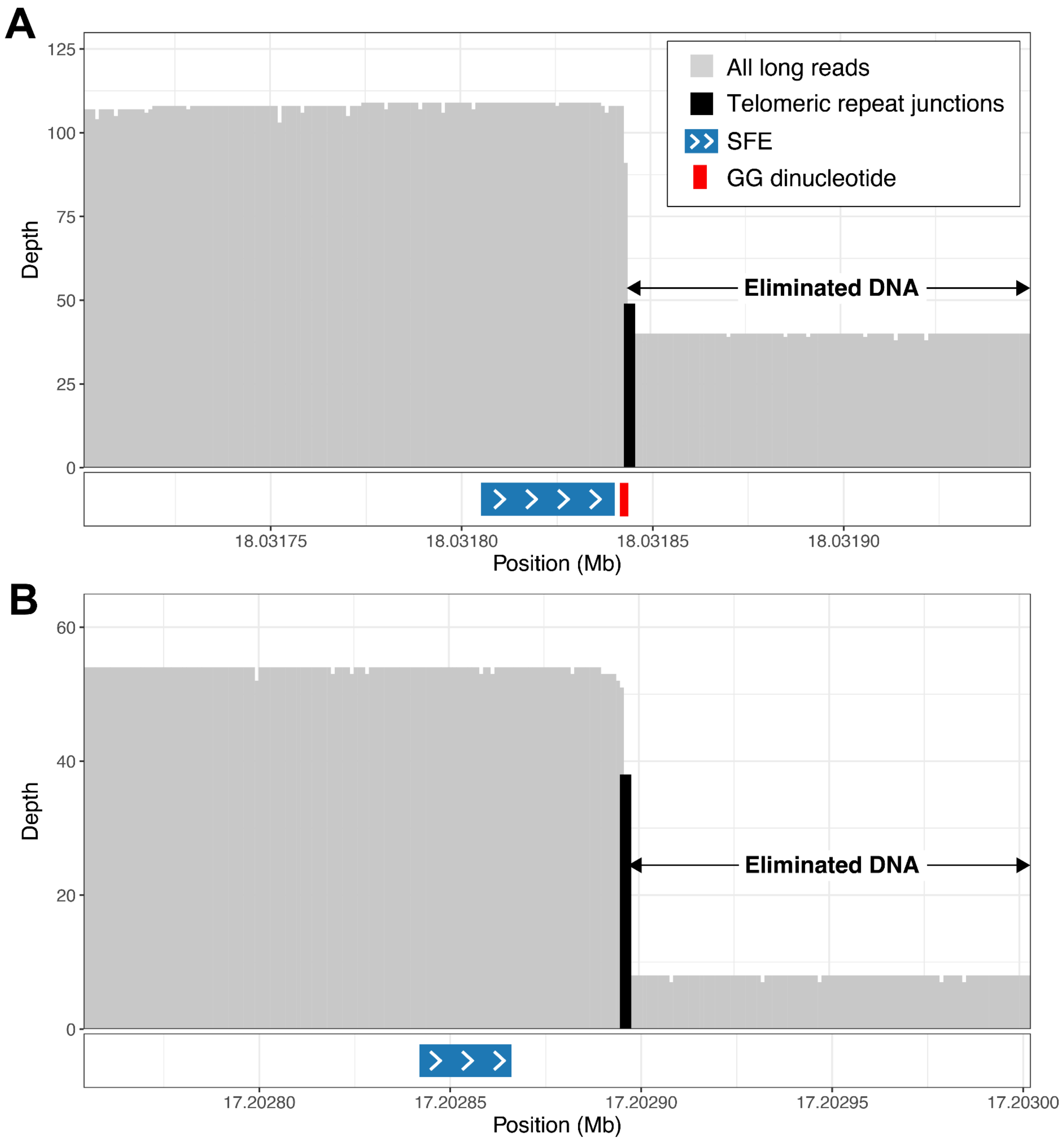


**Fig. S5: Position of the Sequences For Elimination relative to break sites in *C. auriculariae* and *C. parvicauda***

The position of the Sequences for Elimination (SFEs) relative to the break site at (A) the right end of I in *C. auriculariae* (I:18,031,700-18,031,950) and (B) the right end of II in *C. parvicauda* (II:17,202,753-17,203,003). Depth of all PacBio HiFi reads and of junctions between unique sequence and soft-clipped telomeric repeat are shown.

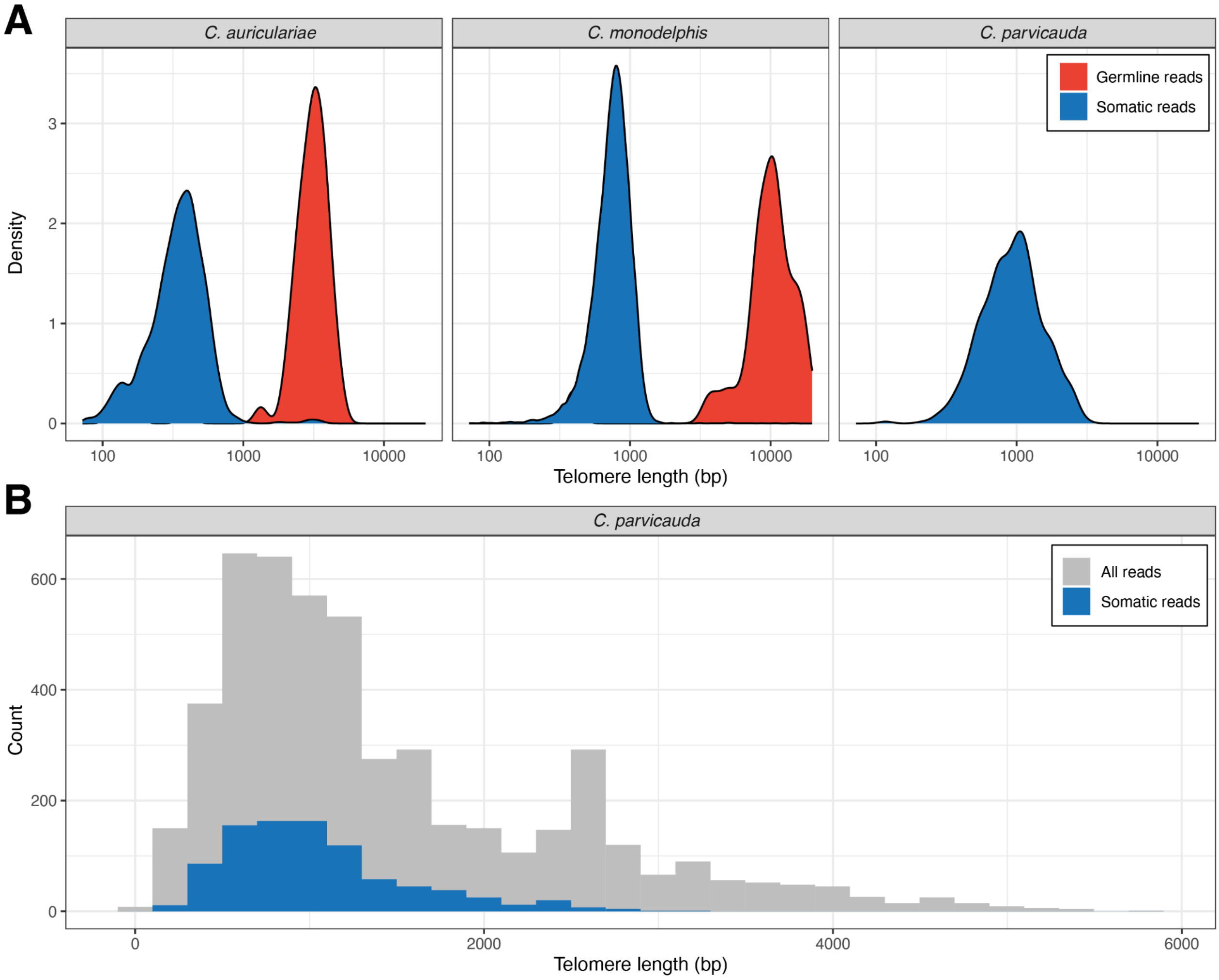


**Fig. S6: Somatic telomeres are shorter than germline telomeres**
(A) Somatic and germline telomere lengths in *C. auriculariae*, *C. monodelphis*, and *C. parvicauda*. Somatic telomere-containing reads were defined as those that aligned to regions 100 bp upstream of each elimination site that contained at least ten consecutive TTAGGC repeats. Germline telomere-containing reads were defined as those that aligned to the first or last 100 bp of each chromosome that contained at least ten consecutive TTAGGC repeats. Germline telomere-containing reads could not be identified in *C. parvicauda* because the germline chromosome ends are not resolved in our assembly. (B) Telomere lengths in somatic telomere-containing reads and in all reads containing telomeric repeats arrays in *C. parvicauda*. A small number of reads with very long repeat arrays (>4 kb) suggest that germline telomeres are also longer than somatic telomeres in *C. parvicauda*.

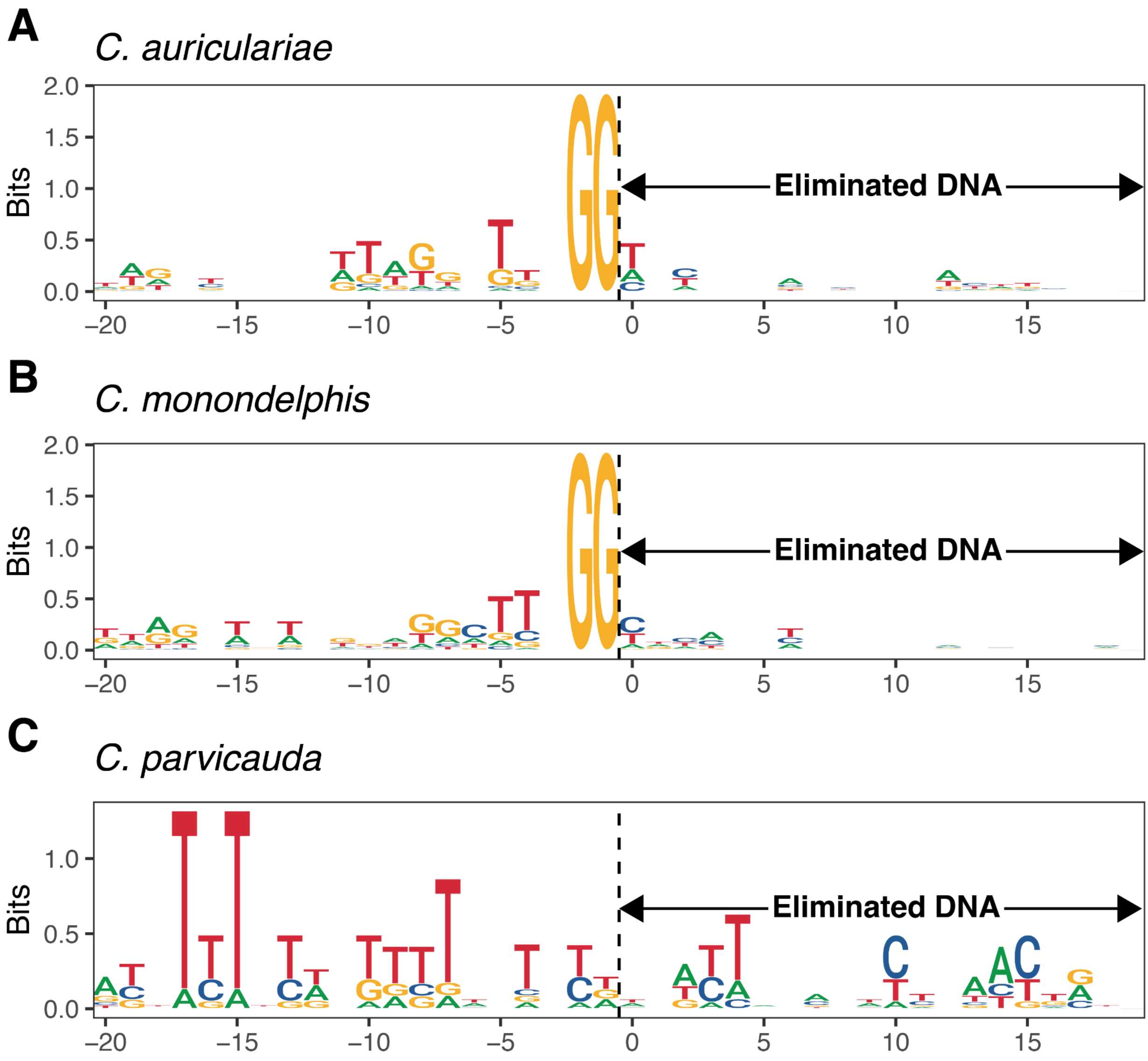


**Fig. S7: Telomere addition sites are preceded by 'GG' dinucleotides in *C. auriculariae* and *C. monodelphis***

Sequence logos of the regions surrounding each telomere addition site in (A) *C. auriculariae,* (B) *C. monodelphis*, and (C) *C. parvicauda*. The two variable telomere addition sites in *C. monodelphis* were excluded.

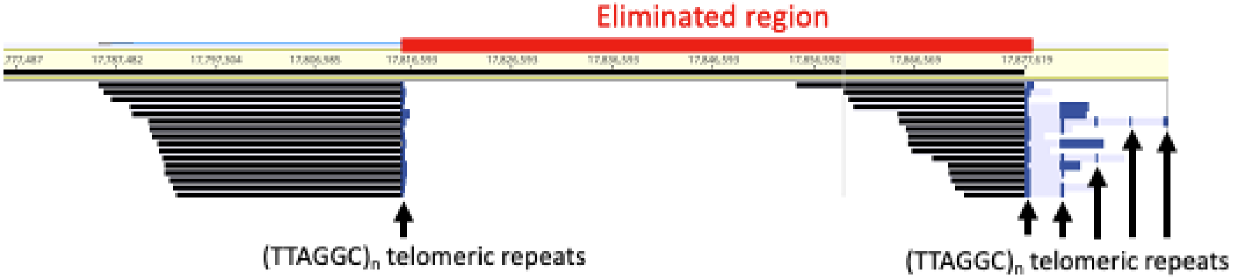


**Fig. S8: Alternative lengthening of telomeres (ALT) in *C. auriculariae***

PacBio HiFi reads (left) aligned to the somatic right end of chromosome II terminate in simple telomeric repeats (dark blue), whereas reads mapped to the germline chromosome ends show ALT patterns in which telomeric repeats (dark blue) are separated by long, non-telomeric sequences (light gray).

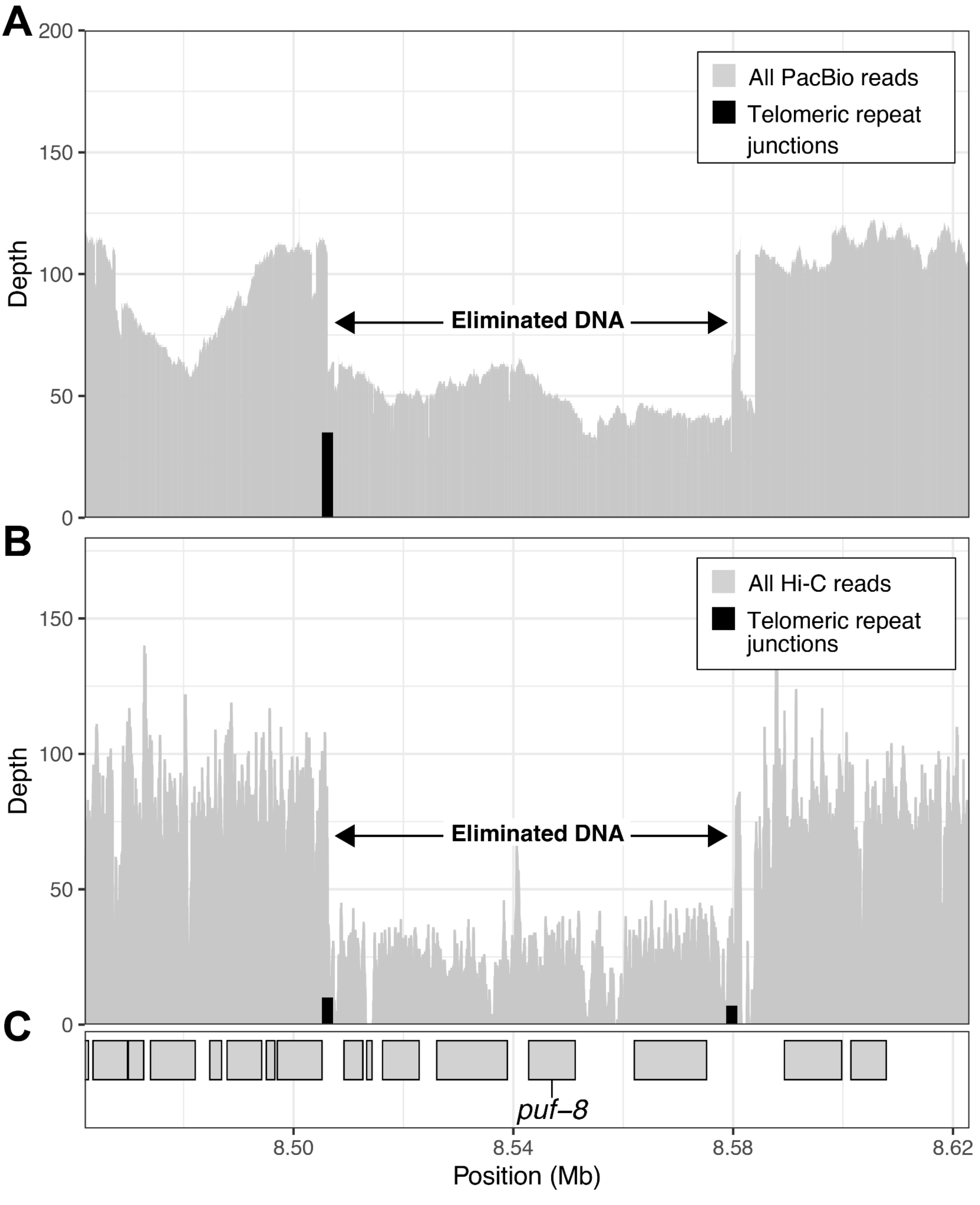


**Fig. S9: Eliminated region containing the *C. auriculariae* orthologue of *puf-8***
(A) Depth of all PacBio HiFi reads and of junctions between unique sequence and soft-clipped telomeric repeat across an eliminated region on chromosome V containing *puf-8* (V: 8.462-8.623 Mb). Only reads containing a telomeric repeat array size of 10 or greater are shown. (B) Depth of all Hi-C reads and of junctions between unique sequence and soft-clipped telomeric repeat across an eliminated region on chromosome V containing *puf-8* (V: 8.462-8.623 Mb). Only reads containing a telomeric repeat array size of 3 or greater are shown. (C) Position of protein-coding genes. *puf-8* is highlighted.

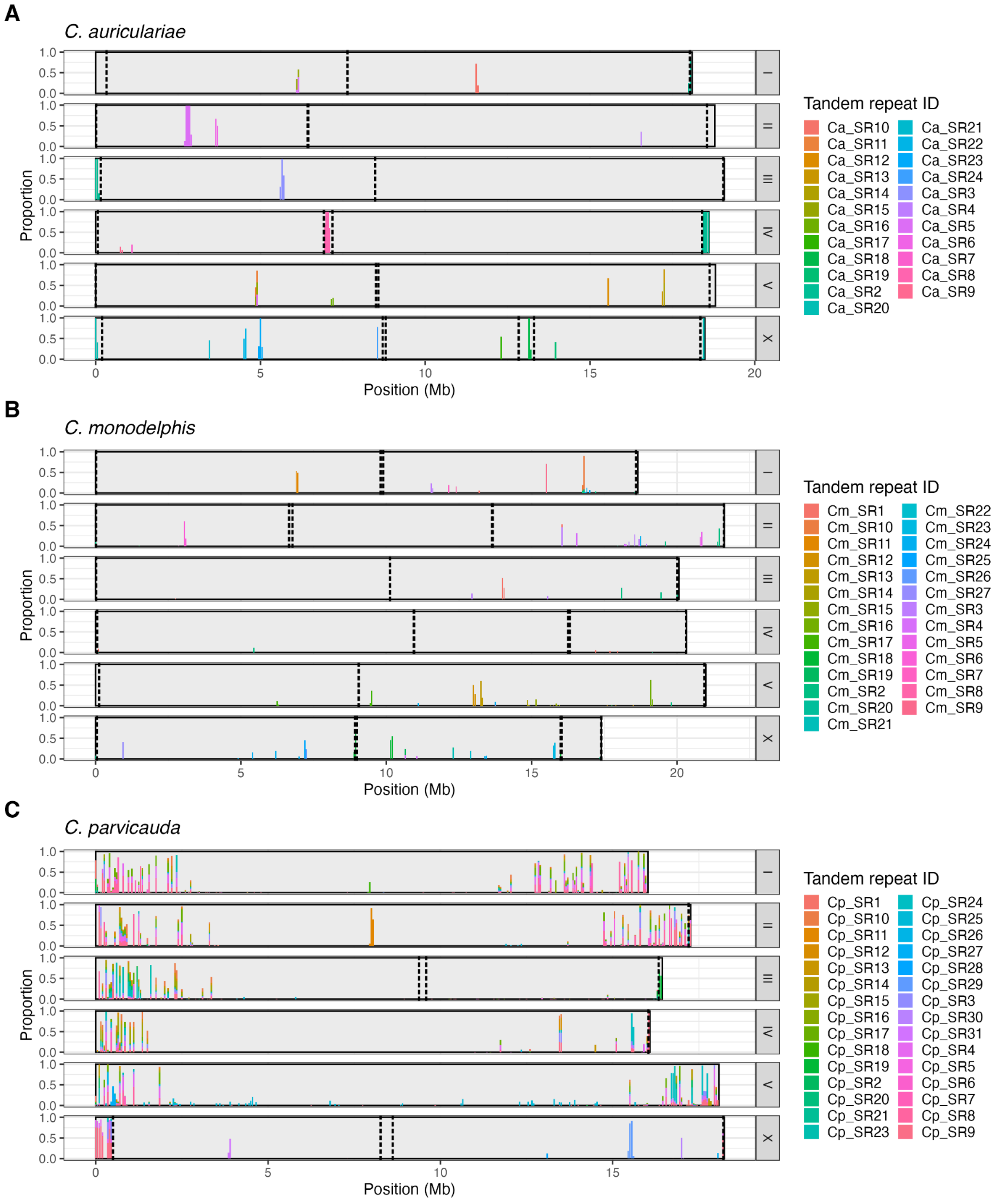


**Fig. S10: Satellite repeats in the *C. auriculariae*, *C. monodelphis*, and *C. parvicauda* genomes**

Locations of satellite repeats in 50 kb windows of the (A) *C. auriculariae,* (B) *C. monodelphis*, and (C) *C. parvicauda* genomes. Locations of telomere addition sites are shown as vertical dotted lines. We defined satellite repeats as units that were at least 50 bp in length that occurred at least 100 times in the genome with an average nucleotide identity of 85%. Probes were designed against two repeats found exclusively in the eliminated regions in *C. auriculariae* (Ca_SR2 and Ca_SR8) and used in our FISH analysis (Fig. 4; Fig. S10).

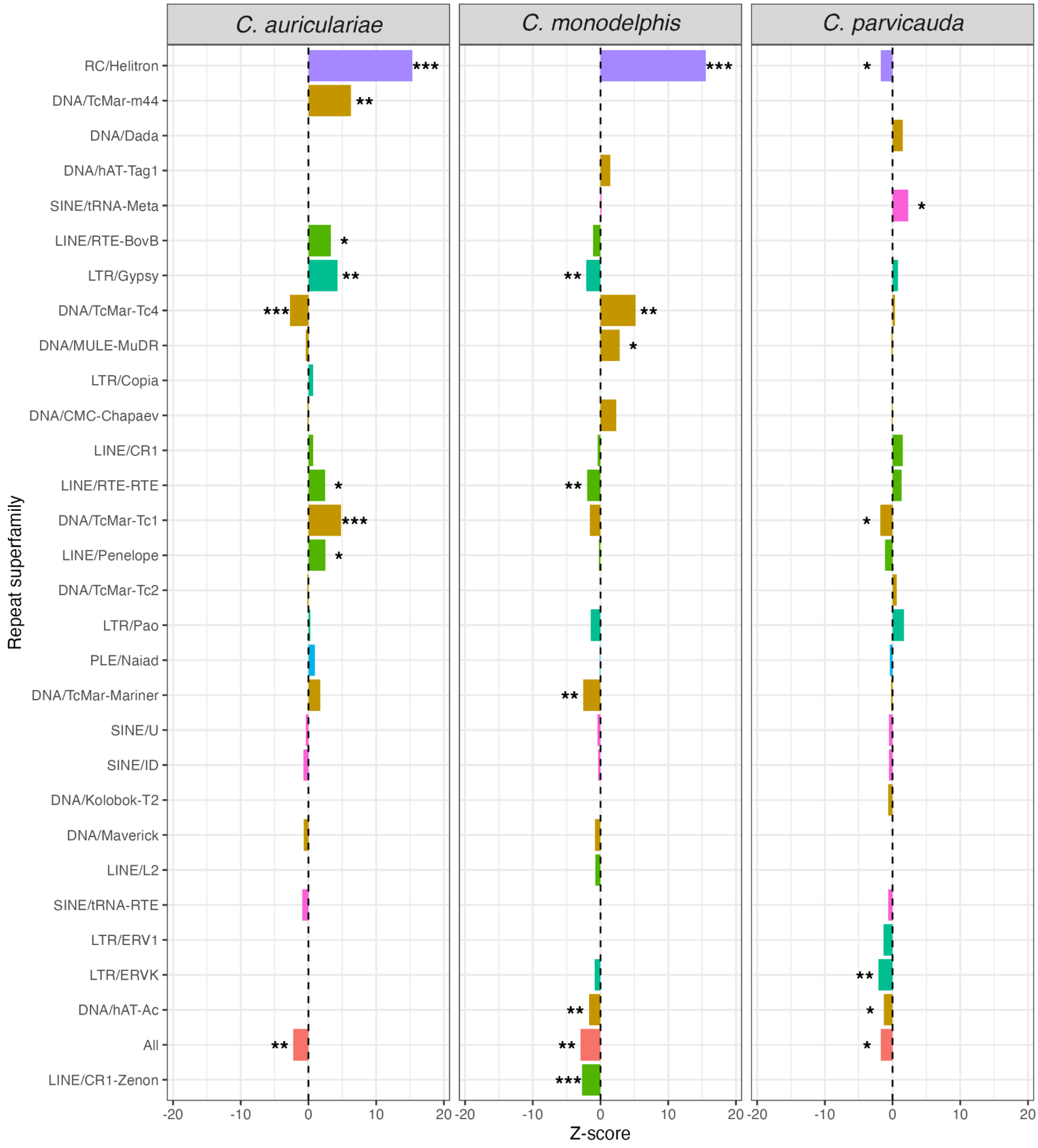


**Fig. S11: Transposable element enrichment in eliminated DNA**

Z-scores of transposable element superfamilies enrichment in DNA that is eliminated from the *C. auriculariae, C. monodelphis*, and *C. parvicauda* genomes. Positive values indicate superfamilies enriched in eliminated DNA, while negative values indicate superfamilies that are significantly depleted. Transposable element annotations are derived from Earl Grey. Only superfamilies with a Z-score ≥ 0.5 or ≤ –0.5 in at least one of the three species are shown. Asterisks indicate statistically significant enrichment (*** = $P < 0.001$, ** = $P < 0.01$, * = $P < 0.05$).

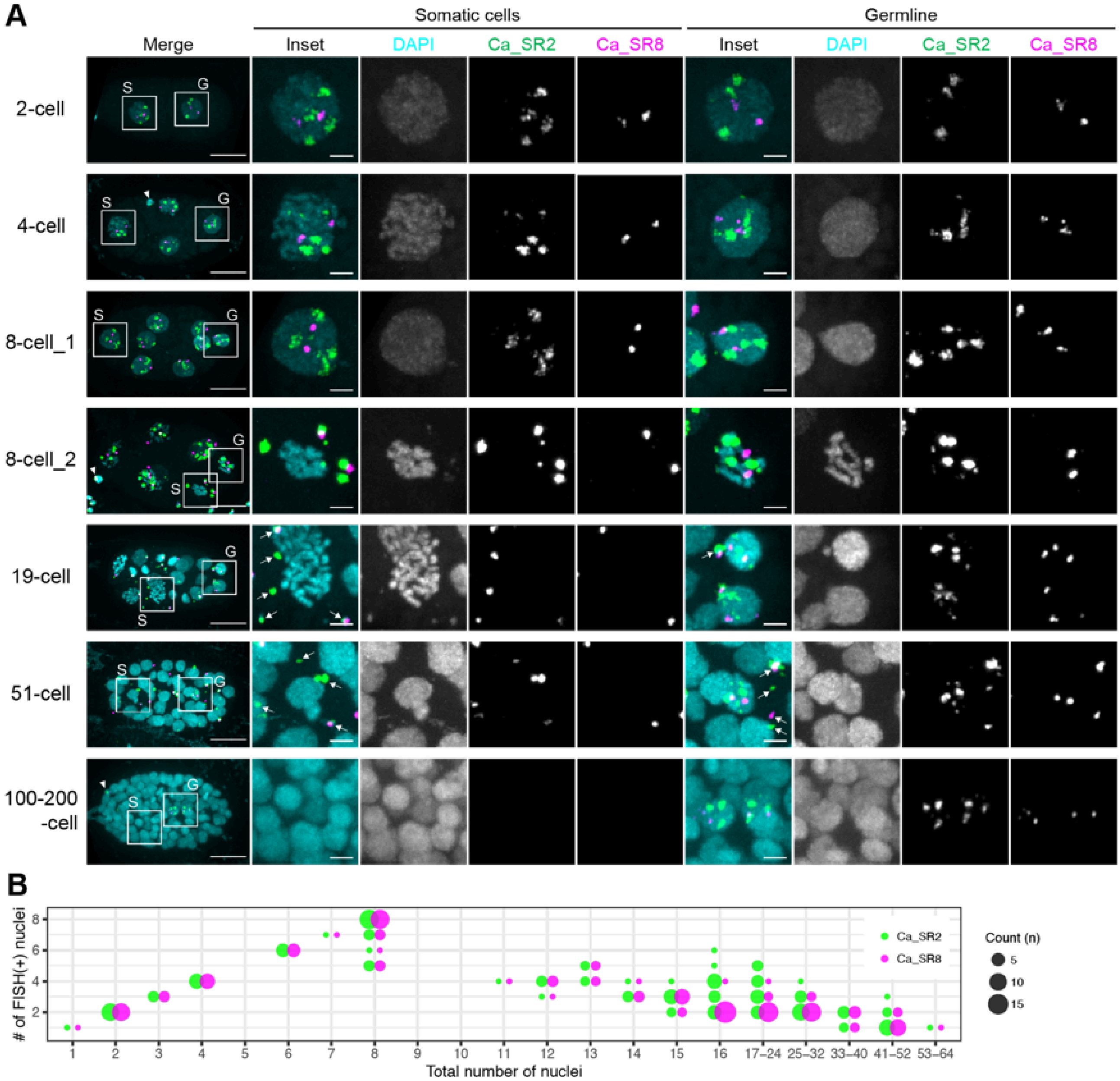


**Fig. S12: FISH analysis of eliminated satellite repeats in *C. auriculariae* embryos**

(A) FISH detection of Ca_SR2 and Ca_SR8 satellite repeats. Insets show magnified views of boxed regions. S, somatic cells; G, germline. Arrows, cytoplasmic DNA fragments; arrowheads, polar bodies. Scale bars: 10 μm (embryos), 2 μm (insets). (B) Quantification of nuclei with Ca_SR2 and Ca_SR8 foci (n = 141 embryos). X-axis: total nuclei per embryo (DAPI). Y-axis: nuclei with FISH signals comparable to germline cells. Dot size indicates number of embryos assayed.

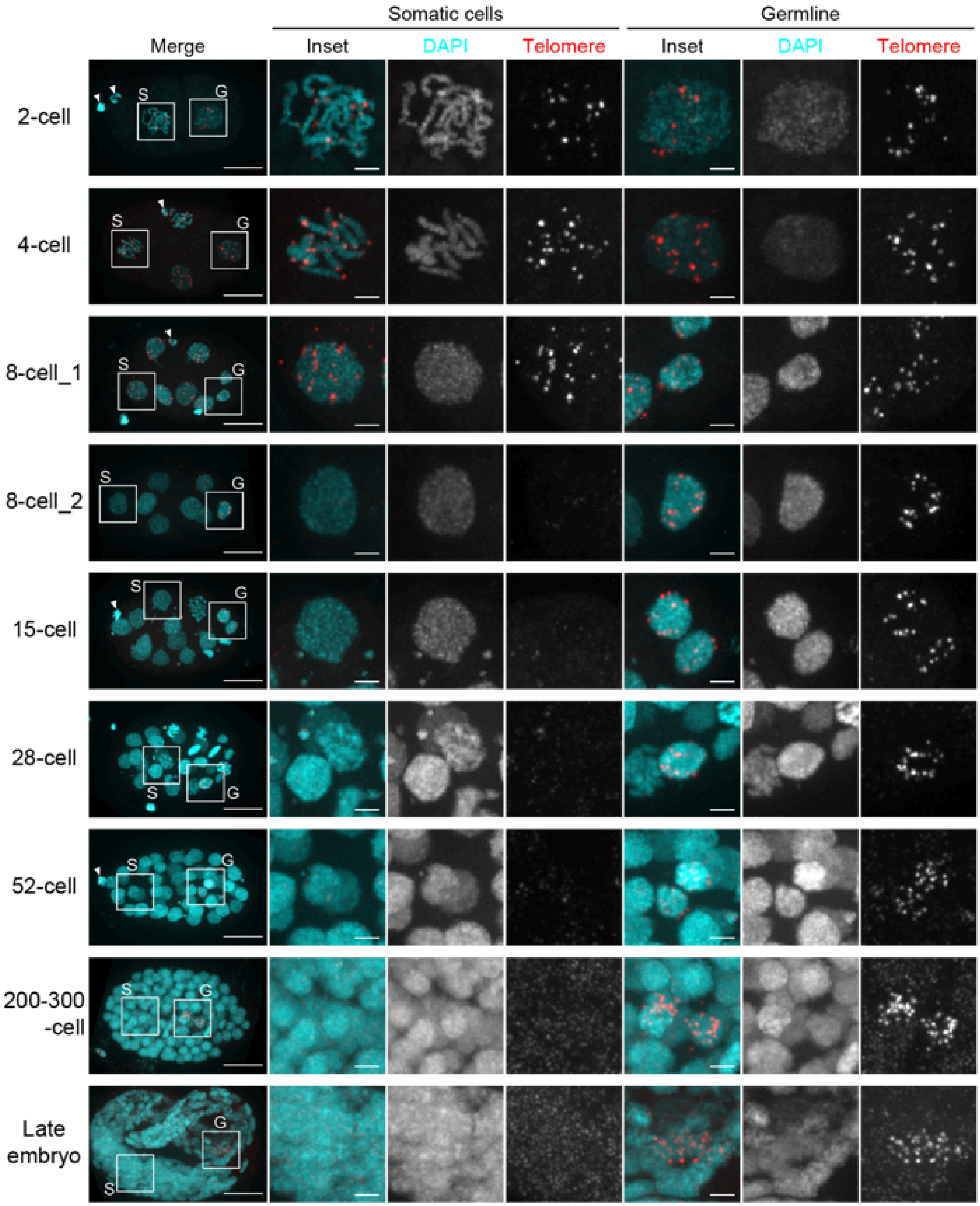


**Fig. S13: FISH analysis of telomeric repeats in *C. auriculariae* embryos**

FISH detection of telomeric repeats. Insets show magnified views of boxed regions. S, somatic cells; G, germline. Arrowheads, polar bodies. Scale bars: 10 µm (embryos), 2 µm (insets).

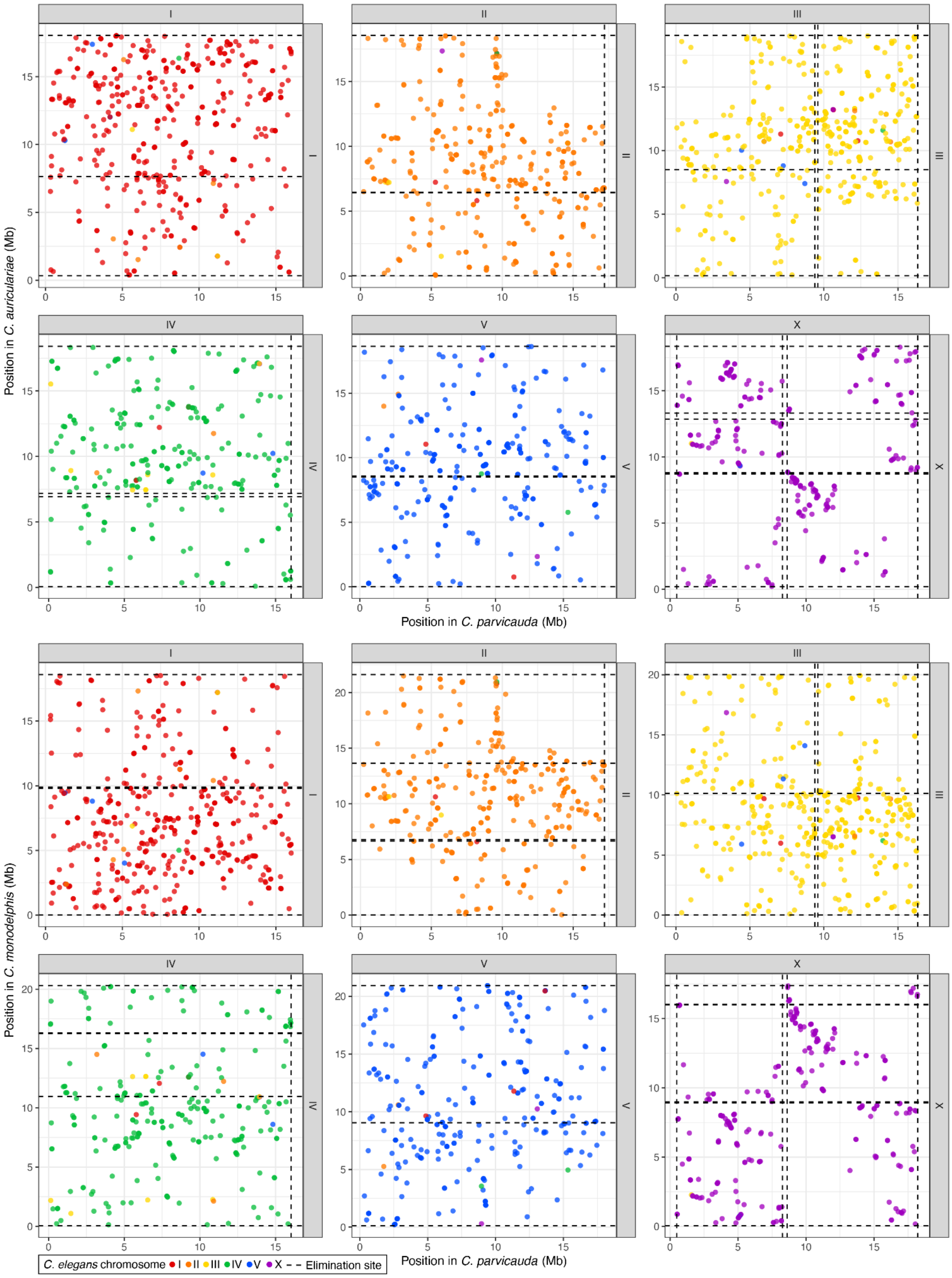

**Fig. S14: Gene order comparison between *C. parvicauda, C. auriculariae* and *C. monodelphis***

Oxford plot showing the relative position of (A) 2,100 BUSCO genes in the *C. parvicauda* and *C. auriculariae* genomes and (B) 2,190 BUSCO genes in the *C. parvicauda* and *C. monodelphis* genomes, coloured by the C. elegans chromosome (I-V, X) in which the orthologous gene is found. Genes that are on different chromosomes or that are not assigned to *C. elegans* chromosomes are not shown. The locations of telomere addition sites are shown with dotted lines.

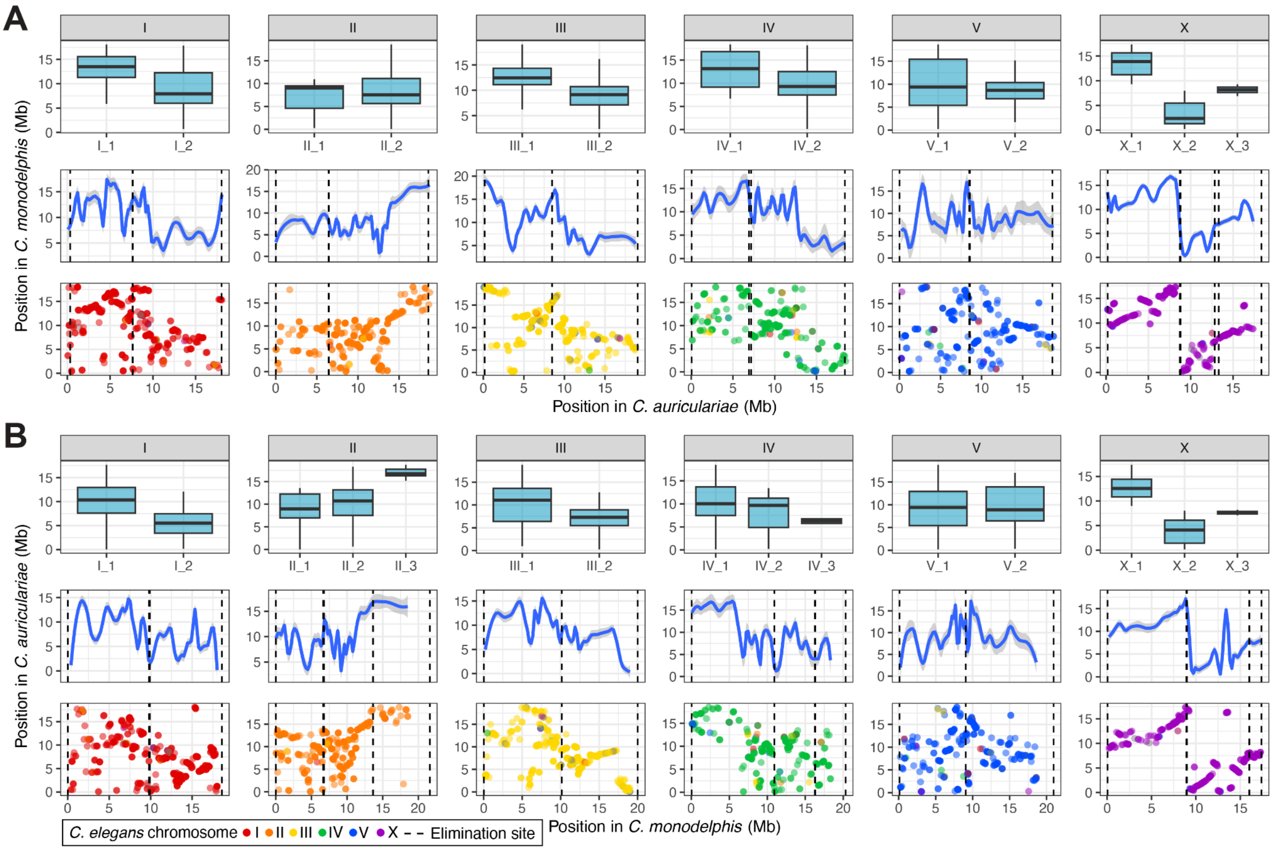


**Fig. S15: Elimination sites sometimes occur between rearrangement domains**

(A) *Caenorhabditis auriculariae* as the focal species, *C. monodelphis* as the comparator. (B) *C. monodelphis* as the focal species, *C. auriculariae* as the comparator. Each column represents one germline chromosome from the focal species. Top row: Boxplots show the genomic positions of BUSCO genes in the comparator species that correspond to each somatic chromosome in the focal species. Middle row: LOESS-smoothed curves represent the average genomic positions of BUSCO genes in the comparator species along the chromosomes of the focal species. Grey shading indicates the standard error. Bottom row: Oxford plots showing the relative positions of BUSCO genes between the focal and comparator species. Vertical dashed lines indicate the locations of elimination sites in the focal species. BUSCO genes are coloured by the C. elegans chromosome (I-V, X) in which the orthologous gene is found

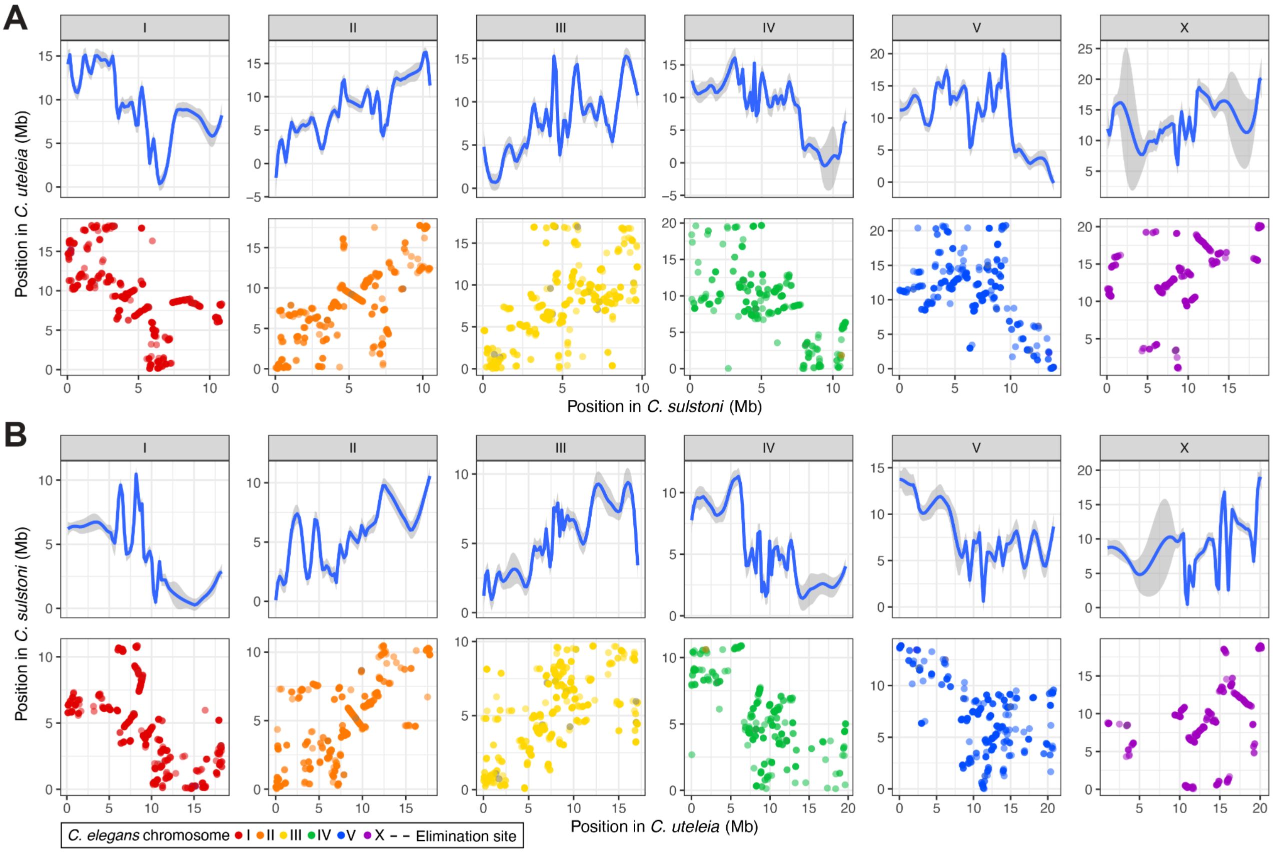


**Fig. S16: Gene order comparison between *C. sulstoni* and *C. uteleia***

(A) *Caenorhabditis sulstoni* as the focal species, *C. uteleia* as the comparator. (B) *C. uteleia* as the focal species, *C. sulstoni* as the comparator. Each column represents one germline chromosome from the focal species. *C. uteleia* and *C. sulstoni* are a similar phylogenetic distance from *C. auriculariae* and *C. monodelphis* (branch length of 0.882 amino acid substitutions per site vs 0.883). Rearrangement domains are apparent in chromosomes IV and V. Top row: LOESS-smoothed curves represent the average genomic positions of BUSCO genes in the comparator species along the chromosomes of the focal species. Grey shading indicates the standard error. Bottom row: Oxford plots showing the relative positions of BUSCO genes between the focal and comparator species. BUSCO genes are coloured by the C. elegans chromosome (I-V, X) in which the orthologous gene is found

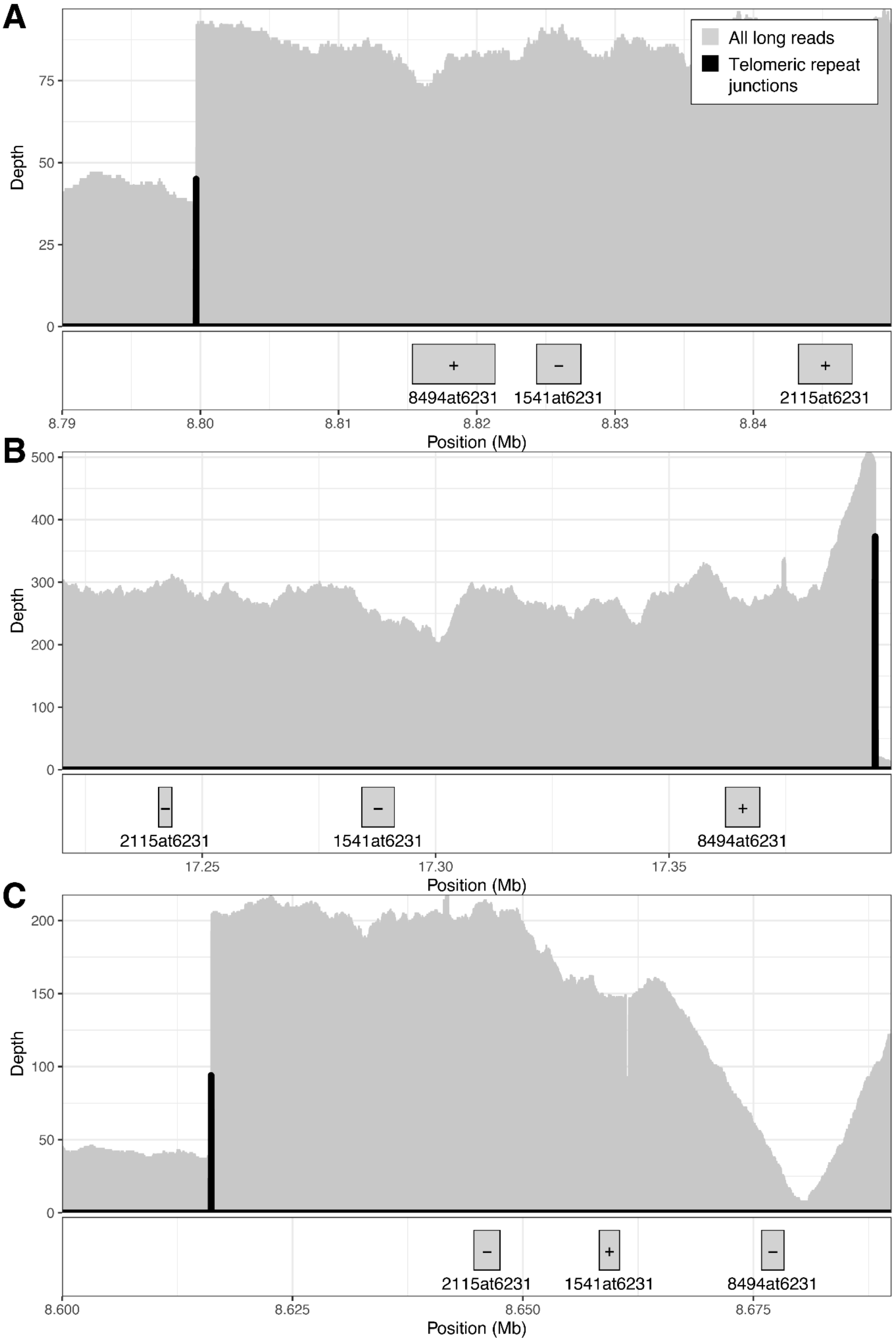


**Fig. S17: Orthologous telomere addition site on X chromosomes of *C. auriculariae, C. monodelphis,* and *C. parvicauda***

Depth of all PacBio HiFi reads and of junctions between unique sequence and soft-clipped telomeric repeat, and the coordinates, IDs, and orientations of upstream BUSCO genes in (A) *C. auriculariae* (X: 8,790,000-8,850,000), (B) *C. monodelphis* (X: 17,220,000-17,398,673), and (C) *C. parvicauda* (X: 8,600,000-8,690,000). '+' indicates that the gene is on the positive strand, whereas '-' indicates that the gene is on the negative strand. The coordinates and orientation of 1541at6231 in *C. monodelphis* were obtained from a second BUSCO run using Augustus as the gene predictor (specified with the --augustus parameter) because the gene was labeled "Missing" in the original BUSCO run, which used MetaEuk as the gene predictor.

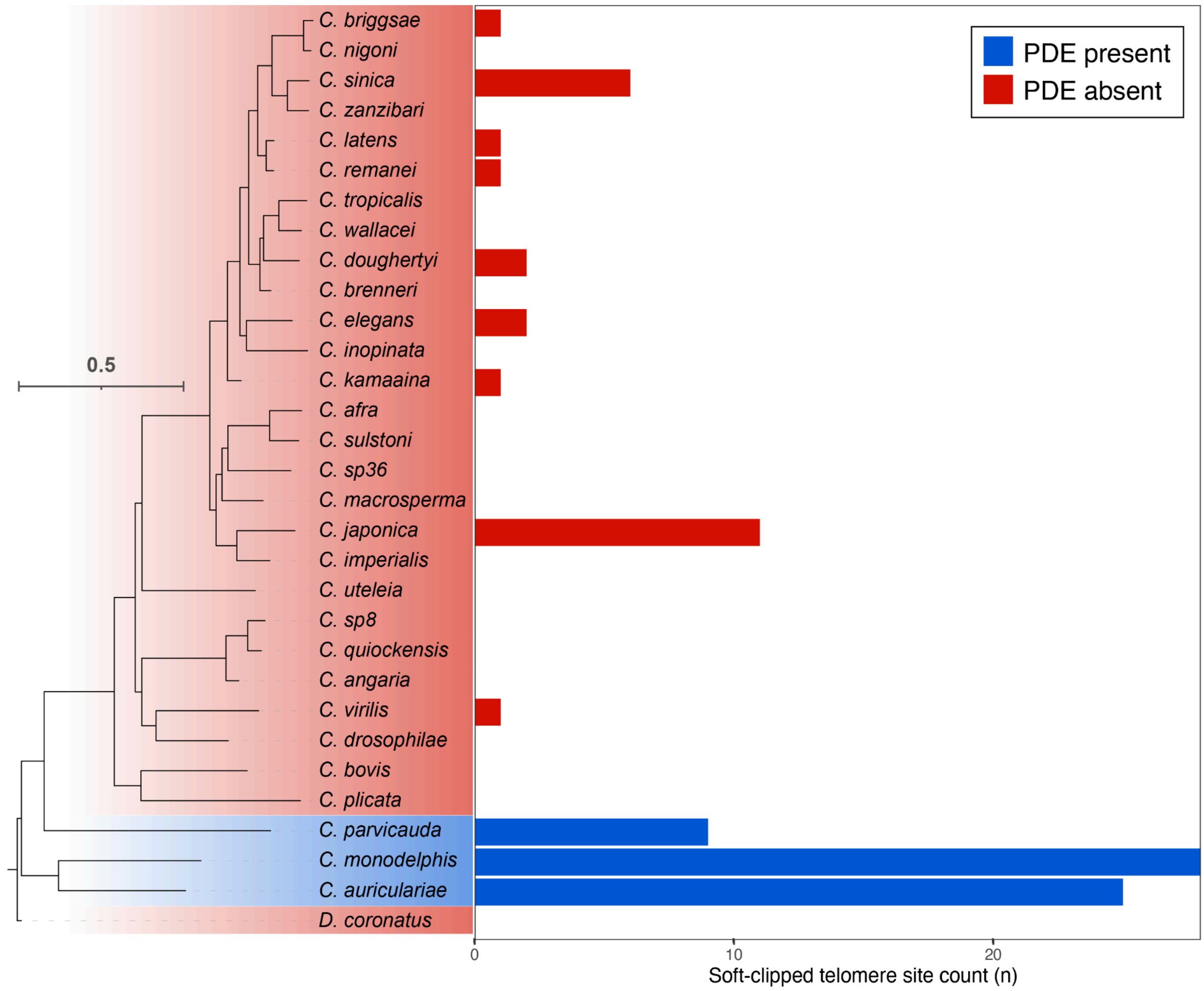


**Fig. S18: Counts of reads containing soft-clipped telomeric repeat**

Bars adjacent to each species indicate the total number of soft-clipped telomere read positions (S2G) identified from long-read alignments using delfies. Sites that were supported by less than 25% of the species' median read coverage were excluded.

#### Table S1: Germline and somatic chromosome counts

| Species | Strain | Germline haploid chromosome count | Somatic haploid chromosome count | Assembly name | Assembly accession |
|---|---|---|---|---|---|
| *C. afra* | JU1286 | 6 | 6 | nxCaeAfra1.1 | GCA_963570955.1 |
| *C. angaria* | PS1010 | 6 | 6 | nxCaeAnga2.1, CAMPv2 | GCA_964213915.1, GCA_947459285.1 |
| *C. auriculariae* | NKZ352 | 6 | 13 | CAUJv3 | GCA_904845305.2 |
| *C. doughertyi* | JU1771 | 6 | 6 | nxCaeDoug1.1 | GCA_963572265.1 |
| *C. drosophilae* | DF5112 | 6 | 6 | nxCaeDros1.1 | GCA_963572285.1 |
| *C. imperialis* | EG5942 | 6 | 6 | nxCaeImpe1.1 | GCA_963572205.1 |
| *C. inopinata* | NKZ35 | 6 | 6 | sp34_v7 | GCA_003052745.1 |
| *C. japonica* | DF5081 | 6 | 6 | nxCaeJapo1.1, CJPJv2 | GCA_963572235.1, (awaiting accession) |
| *C. kamaaina* | QG2077 | 6 | 6 | nxCaeKama2.1 | GCA_964211945.1 |
| *C. macrosperma* | JU2083 | 6 | 6 | nxCaeMacr1.1 | GCA_963932285.1 |
| *C. monodelphis* | JU1667 | 6 | 15 | nxCaeMono1.1 | GCA_964197825.1 |
| *C. niphades* | NKZ392 | 6 | 6 | CNIPv6 | GCA_946814055.1 |
| *C. parvicauda* | NIC534 | 6 | 8 | nxCaeParv1.1 | GCA_963978915.1 |
| *C. plicata* | SB355 | 6 | 6 | nxCaePlic1.1 | GCA_963931815.1 |
| *C. quiockensis* | JU2809 | 6 | 6 | nxCaeQuio2.1 | GCA_964198105.1 |
| *C. sinica* | JU800 | 6 | 6 | nxCaeSini1.1 | GCA_963932045.1 |
| *C. sp. 8* | DF5173 | 6 | 6 | nxCaeSpee1.1 | GCA_963572245.1 |
| *C. sulstoni* | JU2788 | 6 | 6 | nxCaeSuls1.1 | GCA_963966605.1 |
| *C. uteleia* | JU2585 | 6 | 6 | nxCaeUtel1.1 | GCA_963573275.1 |
| *C. virilis* | JU1968 | 6 | 6 | nxCaeViri1.1 | GCA_964036255.1 |
| *C. wallacei* | JU1904 | 6 | 6 | nxCaeWall1.1 | GCA_963932035.1 |
| *C. zanzibari* | JU2190 | 6 | 6 | nxCaeZanz1.1 | GCA_963966625.1 |
| *Diploscapter coronatus* | PDL0010 | 1 | 1 | nxDipCoro1.1 | GCA_964036155.1 |

#### Table S2: Reference genome metrics

| | | | |
|---|---|---|---|
| **Accession information** | | | |
| Species | *Caenorhabditis auriculariae* | *Caenorhabditis monodelphis* | *Caenorhabditis parvicauda* |
| Strain | NKZ352 | JU1667 | NIC534 |
| Version | CAUJv3 | nxCaeMono1.1 | nxCaeParv1.1 |
| Assembly accession | GCA_904845305.2 | GCA_964197825.1 | GCA_963978915.1 |
| **Genome assembly metrics** | | | |
| Span (Mb) | 111.9 | 119.4 | 115.4 |
| Scaffolds (n) | 6 (+MT) | 13 (+MT) | 498 (+MT) |
| Scaffold N50 (Mb) | 18.8 | 20.3 | 16.4 |
| Assembly in six chromosomes (%) | 100 | 99.6 | 88.5 |
| Contigs (n)* | 35 | 38 | 1422 |
| Contig N50 (Mb) | 14.5 | 14.2 | 0.2 |
| Gaps (n) | 29 | 25 | 924 |
| Genome BUSCO completeness (%)† | 95.3 | 93.9 | 89.6 |
| Genome BUSCO duplication (%)† | 1 | 1 | 1.6 |
| **Protein-coding gene metrics** | | | |
| Genes (n) | 15,814 | 16,482 | 15,601 |
| Transcripts (n) | 18,759 | 19,632 | 18,059 |
| Gene set BUSCO completeness (%)† | 97.2 | 97 | 91.6 |

*Contig values were calculated by splitting scaffolds at ≥10 consecutive Ns.
†Genome and gene set completeness was assessed using BUSCO (version 5.2.2) with the nematoda_odb10 dataset (using the Augustus option when assessing genome completeness).

#### Table S3: Eliminated DNA summary

| Chromosome | Size (Mb) | External eliminated regions (n) | Internal eliminated regions (n) | Total eliminated DNA (kb) | Total eliminated DNA (%) |
|---|---|---|---|---|---|
| *C. auriculariae* nxCaeAuri1.1 | | | | | |
| I | 18.1 | 2 | 1 | 401.3 | 2.2 |
| II | 18.8 | 2 | 1 | 276.7 | 1.5 |
| III | 19.1 | 2 | 1 | 183.5 | 1 |
| IV | 18.6 | 2 | 1 | 525.7 | 2.8 |
| V | 18.8 | 2 | 1 | 263.4 | 1.3 |
| X | 18.5 | 2 | 2 | 880.1 | 4.8 |
| Entire genome | 111.9 | 12 | 7 | 2,530.60 | 2.3 |
| *C. monodelphis* nxCaeMono1.1 | | | | | |
| I | 18.7 | 2 | 1 | 171.5 | 0.9 |
| II | 21.6 | 2 | 2 | 170.2 | 0.8 |
| III | 20.1 | 2 | 1 | 74.2 | 0.4 |
| IV | 20.3 | 2 | 2 | 142.7 | 0.7 |
| V | 21 | 2 | 1 | 172.2 | 0.8 |
| X | 17.4 | 2 | 2 | 149.8 | 0.9 |
| Entire genome | 119.4 | 12 | 9 | 880.5 | 0.7 |
| *C. parvicauda* nxCaeParv1.1 | | | | | |
| I | 16 | 0* | 0 | 0 | 0.0 |
| II | 17.3 | 1* | 0 | 60.5 | 0.3 |
| III | 16.4 | 1* | 1 | 312.6 | 1.9 |
| IV | 16.1 | 1* | 0 | 46.8 | 0.3 |
| V | 18.1 | 0* | 0 | 0 | 0.0 |
| X | 18.2 | 2 | 2 | 877.2 | 4.8 |
| SCAFFOLD_210 | 0.02 | 1 | 0 | 15.8 | 79.0 |
| Entire genome | 115.4 | 6* | 4 | 1,313.00 | 1.1 |

*We do not know if elimination occurs at the remaining chromosome ends in *C. parvicauda* as they are unresolved in our assembly.

#### Table S4: Enrichment of satellite repeat in eliminated DNA

| | *C. auriculariae* | *C. monodelphis* | *C. parvicauda* |
|---|---|---|---|
| Genome span (Mb) | 111.86 | 119.43 | 115.36 |
| Satellite repeat span (Mb) | 1.62 | 1.22 | 24.80 |
| % satellite repeat | 1.45% | 1.02% | 21.50% |
| Eliminated span (Mb) | 2.53 | 0.88 | 1.31 |
| % of eliminated DNA that is satellite repeat | 25.83% | 8.50% | 38.72% |
| Expected eliminated satellite repeat span (Mb) | 0.04 | 0.01 | 0.28 |
| Observed eliminated satellite repeat span (Mb) | 0.65 | 0.07 | 0.51 |
| Fold enrichment | 17.87 | 8.32 | 1.80 |
| % of satellite repeat that is eliminated | 40.42% | 6.13% | 2.05% |
| P-value (from regioneR) | 0.00 | 0.02 | 0.00 |

#### Table S5: Orthologous telomere addition sites

| Species 1 | | | Species 2 | | | | |
|---|---|---|---|---|---|---|---|
| **Species** | **Chromosome** | **Telomere addition site** | **Species** | **Chromosome** | **Telomere addition site** | **Overlapping BUSCOs (n)** | **Overlapping BUSCO IDs** |
| *C. auriculariae* | I | 7642713 | *C. monodelphis* | I | 18589543 | 2 | 3712at6231,9060at6231 |
| *C. auriculariae* | III | 19044455 | *C. monodelphis* | III | 19931 | 2 | 3084at6231,5501at6231 |
| *C. auriculariae* | X | 12835912 | *C. monodelphis* | X | 46284 | 2 | 2037at6231,3574at6231 |
| *C. auriculariae* | X | 13299083 | *C. monodelphis* | X | 16032226 | 2 | 1927at6231,6438at6231 |
| *C. auriculariae* | X | 8708168 | *C. monodelphis* | X | 15997345 | 2 | 3388at6231,5755at6231 |
| *C. auriculariae* | X | 8799697 | *C. monodelphis* | X | 17394178 | 3 | 2115at6231,1541at6231,8494at6231* |
| *C. auriculariae* | X | 8799697 | *C. parvicauda* | X | 8616169 | 3 | 1541at6231,2115at6231,8494at6231 |
| *C. monodelphis* | X | 17394178 | *C. parvicauda* | X | 8616169 | 3 | 2115at6231,1541at6231,8494at6231* |

*The coordinates and orientation of 1541at6231 in C. monodelphis were obtained from a second BUSCO run using Augustus as the gene predictor (specified with the --augustus parameter) because the gene was labeled “Missing” in the original BUSCO run, which used MetaEuk as the gene predictor.

#### Table S6: Gene prediction metrics and accessions

| Species | Accession | RNA-seq source accession | Genes (n) | Proteins (n) | BUSCO completeness (%) |
|---|---|---|---|---|---|
| *C. afra* | GCA_963570955.1 | ERR1059227 | 19096 | 21765 | 98.9 |
| *C. angaria* | GCA_964213925.1 | ERR13319081 | 17801 | 19898 | 97.8 |
| *C. auriculariae* | GCA_904845305.2 | DRR252169 | 15814 | 18759 | 97.4 |
| *C. doughertyi* | GCA_963572265.1 | ERR1039279 | 28351 | 32554 | 99.7 |
| *C. drosophilae* | GCA_963572285.1 | ERR15946064 | 12924 | 15037 | 98.2 |
| *C. imperialis* | GCA_963572205.1 | ERR13319091 | 19390 | 22477 | 98 |
| *C. japonica* | GCA_963572235.1 | ERR13319090 | 20639 | 24236 | 98.6 |
| *C. kamaaina* | GCA_964211945.1 | ERR13362831 | 20508 | 23322 | 99.3 |
| *C. latens* | GCA_002259235.3 | SRR5831583 | 25520 | 28534 | 99.5 |
| *C. macrosperma* | GCA_963932285.1 | ERR1018632,ERR1018631,ERR1018630 | 23804 | 27690 | 99.3 |
| *C. monodelphis* | GCA_964197825.1 | ERR690851 | 16482 | 19632 | 97 |
| *C. parvicauda* | GCA_963978915.1 | ERR1233391,ERR1233392 | 15601 | 18059 | 91.6 |
| *C. plicata* | GCA_963931815.1 | ERR1059194 | 16420 | 18798 | 95.3 |
| *C. quiockensis* | GCA_964198105.1 | ERR1055246 | 19733 | 22202 | 98.2 |
| *C. sinica* | GCA_963932045.1 | ERR15954981 | 27205 | 31987 | 98.3 |
| *C.* sp. 8 | GCA_963572245.1 | ERR15954974 | 15865 | 18038 | 98.1 |
| *C. sulstoni* | GCA_963966605.1 | ERR1233611 | 17926 | 20574 | 98.7 |
| *C. tropicalis* | GCA_043792875.1 | SRR31543810 | 20708 | 23634 | 99.3 |
| *C. uteleia* | GCA_963573275.1 | ERR1233605,ERR1233606 | 26204 | 30204 | 98.6 |
| *C. virilis* | GCA_964036255.1 | ERR1055674 | 18157 | 20799 | 98.3 |
| *C. wallacei* | GCA_963932035.1 | ERR690202 | 20358 | 23038 | 99.2 |
| *C. zanzibari* | GCA_963966625.1 | ERR1233498 | 22434 | 25409 | 99.4 |
| *Diploscapter coronatus* | GCA_964036155.1 | ERR13319096 | 29547 | 34431 | 93.9 |

### Supplementary Text

#### Extension of *C. monodelphis* chromosome ends

##### Chromosome I

The original left end of chromosome I (IL) had a high identity alignment spanning approximately 20 kb to a contig (ptg000050l) from the all-read hifiasm assembly that terminated in a ~10 kb telomere repeat array. We used CAP3 (with a minimum overlap length of 10 kb) to join ptg000050l to IL.

The original IR contained an inverted tandem repeat structure, which was not supported by any aligned reads. We manually removed the sequence after the inversion junction (18,616,122 bp) using the getfasta function from BEDtools. We identified a ~38 kb contig in the telomere-only hifiasm assembly that shared the same repeat structure and that terminated in a ~10 kb telomere repeat array. We used agptools to scaffold this contig onto the IR (separated by a 200 bp gap).

##### Chromosome II

The original IIL did not terminate in a telomere repeat array. Both reads and aligned contigs terminated in a low complexity repeat that was absent from the original IIL. We identified a ~14 kb PacBio HiFi read (m64174e_210924_183014/1180280/ccs) that had a ~9 kb alignment to IIL and that contained ~5 kb of the low complexity repeat. We used CAP3 (minimum overlap length of 5 kb) to join this read to IIL.

The original IIR ended in low complexity repeat, but not in a telomere repeat array. We identified a ~11 kb read (m64174e_210924_183014/72747242/ccs) that ended in a germline-length (~4 kb) telomere repeat array and had a ~6 kb alignment to IIR. We used CAP3 (minimum overlap length of 2 kb) to join this read to IIR.

##### Chromosome III

We identified a ~15 Mb contig (ptg000004l) from the all-read hifiasm assembly that aligned to IIIL and terminated in a germline-length (~15 kb) telomere repeat array. We extracted the first 50 kb of ptg000004l and used CAP3 (minimum overlap length of 10 kb) to join it to IIIL.

We identified a ~60 kb contig (ptg000041l) from the all-read hifiasm assembly that had a ~12 kb alignment to the original IIIR and that terminated in a germline-length (~14 kb) telomere repeat array. We used CAP3 (minimum overlap length of 10 kb) to join ptg000041l to IIIR.

##### Chromosome IV

IVL did not end in a telomere repeat array, but we could not identify any reads or contigs suitable for extending the chromosome end.

We identified a ~1.2 Mb contig (ptg000024l) from the all-read hifiasm assembly that had a ~34 kb alignment to IVR. We extracted the last 50 kb of ptg000024l and used CAP3 (minimum overlap length of 10 kb) to join to IVR.

##### Chromosome V

The original VL ended in a complex tandem repeat and lacked a telomere repeat array. We extracted all reads that had a primary alignment to the first 34 kb of VL with a map quality of 60 using samtools (view -h -q 60 -F 2048) and assembled them using hifiasm. We identified an ~85 kb contig in this assembly (ptg000001l) that contained a repeat structure identical to that of VL and that ended in a ~8 kb telomere repeat array. We used agptools to add this contig to VL (separated by a 200 bp gap).

We identified a ~40 kb contig (ptg000016l) that had a ~15 kb alignment to VR. We used CAP3 (minimum overlap length of 10 kb) to join ptg000016l to VR.

##### Chromosome X

XL and XR both end in low complexity repeat and we could not identify any reads or contigs suitable for extending the chromosome ends.

The internally eliminated region 16.00 - 16.03 Mb contained a gap, to which the somatic telomere array was assembled. We identified the contig that comprised the somatic telomere array (ptg000011l) and removed the telomere sequence using BEDtools getfasta. We then identified the longest soft-clipped germline read that aligned to this contig (m64174e_210923_123112/43910428/ccs) and used this to extend the contig with CAP3 (minimum overlap length of 5 kb). The scaffold was then reassembled using agptools.

##### Data S1

Details of sequence for elimination (SFEs) in each genome.

##### Data S2

Details of genes eliminated from each genome.

##### Movie S1

Live imaging of a *C. auriculariae* 4-cell embryo expressing GFP::Histone H2B.

##### Movie S2

Live imaging of an 8-cell *C. auriculariae* embryo expressing GFP::Histone H2B.
